# Hematogenous neuroinvasion and genotype-dependent transmission of influenza A H5N1 viruses in the cat host

**DOI:** 10.64898/2026.02.21.707182

**Authors:** Salman L. Butt, Ruchi Rani, Mohammed Nooruzzaman, Elena A. Demeter, Pablo S.B. de Oliveira, Gavin R. Hitchener, Diego G. Diel

## Abstract

The spillover of highly pathogenic avian influenza (HPAI) A H5N1 virus to mammalian hosts raises major concerns due to its pandemic potential. Cats are frequently affected mammals, often succumbing to systemic and neurological disease. Here, we characterized the pathogenesis and transmissibility of two H5N1 genotypes, B3.13 and D1.1, in cats. Infected cats exhibited high-level viremia and virus shedding in nasal, oral, and fecal secretions were consistently detected. The virus replicated initially in the upper respiratory tract and lungs, followed by systemic dissemination and neuroinvasion. Notably, the virus crossed the blood-brain-barrier by infecting endothelial cells, spreading to astrocytes and neurons, causing multifocal encephalitis. D1.1-virus infection caused protracted disease with lower shedding and no transmissibility, whereas B3.13 virus caused rapid onset with efficient shedding and transmission. These findings reveal critical H5N1 neuropathogenesis mechanisms and highlight mammalian transmission potential in a species with close human contact.

## Introduction

Highly pathogenic avian influenza (HPAI) H5N1 clade 2.3.4.4b viruses infect a broad range of host species.^1–4^ In the United States, a newly emerged reassortant genotype, H5N1 clade 2.3.4.4b B3.13, was first detected in 2023 and, since then, has caused disease outbreaks in multiple species.^5–9^ In March 2024, this virus spilled over from avian hosts to dairy cattle, with evidence of sustained cow-to-cow transmission, spillover from cows-to-cats and subsequent spillback from cows-to-birds, highlighting its expanded host range.^5^

As H5N1 clade 2.3.4.4b viruses continue to circulate in wild birds and mammals, new genotypes and variants are continuously emerging^10^. Between 2021-2022, fully Eurasian (Eu) lineage 2.3.4.4b H5N1 viruses of genotypes A1-A5 were introduced to North America.^11^ These viruses reassorted^12^ with endemic low pathogenic avian influenza viruses (LPAI) of American (Am) lineages, leading to multiple descendant genotypes.^3^ The major reassortant genotypes identified include B-type genotypes (derived from A1 genotype), C-type genotypes (from A2 genotype), and D-type genotypes (reassortants of the A3 Eurasian genotype with LPAI Am lineage viruses).^12^ In October 2024, a novel genotype, D1.1 – descendant from the fully Eu A3 genotype – emerged in wild birds in North America^13^ causing outbreaks in poultry and infections in dairy cattle, cats, humans and wild mammalian hosts.^10^ To date, the B3.13 and D1.1 genotypes have proven to be the most successful reassortant H5N1 viruses in North America, causing widespread outbreaks across both wild and domestic bird populations and crossing species barriers to infect mammals.^10^ Despite this expansion in host-range, the differences in viral pathogenesis, replication, and transmission dynamics between these viruses in avian and mammalian hosts remain poorly understood.

Domestic cats (*Felis catus*) are particularly susceptible to H5N1 infection.^5,8,14–16^ Prior to 2022, only 38 feline H5N1 cases had been reported worldwide, with none in North America.^10^ While just one domestic cat infection was documented in the United States between 2022 and 2023, the emergence and spillover of H5N1 genotype B3.13 viruses in dairy cattle in 2024 led to a marked surge, with 266 feline cases reported by January 26, 2026.^10^ The infection often proves fatal in cats^17–19^ causing neurological disease that frequently results in death.^17^ Cats are thought to contract H5N1 virus through ingestion of infected birds or consumption of contaminated raw milk from affected dairy cows.^5^ The high viral loads found in brain tissue correlate with the presence of the influenza sialic acid (SA) receptors (α2,3- and α2,6-linked) in neural tissues, potentially explaining the severe neurological manifestations observed in infected cats.^15,17^ However, the understanding of many aspects of the pathogenesis, infection dynamics, and transmissibility of H5N1 clade 2.3.4.4b viruses in felids remains incomplete.

In this study, we investigated the infection dynamics, tissue tropism, and pathways of neuroinvasion of H5N1 virus in cats. Additionally, we compared the transmission efficiency of a bovine-derived B3.13 virus with that of an avian-derived D1.1 virus to assess genotype-specific differences in transmission potential in this highly relevant susceptible host species.

## Results

### H5N1 genotype B3.13 virus infection caused clinical disease in cats

We assessed the H5N1 B3.13 virus infection dynamics and disease progression in cats following oronasal inoculation. For this, twelve (n = 12) H5N1-seronegative research cats were randomly allocated to two experimental groups: a mock-infected control group (G1, n = 3), and an H5N1-infected group (G2, n = 9, A/Cattle/Texas/063224-24-1/2024; B3.13 [TX2/24]). Animals in G1 were mock inoculated oronasally (1 mL orally, 1 mL nasally [0.5 mL per nostril]) with minimum essential media (MEM), whereas animals in G2 were inoculated through the same route with 2 x 10^4.5^ PFU of B3.13 (TX2/24) virus (Figure 1A). Following inoculation, animals were monitored, and individual clinical scores were recorded daily following a clinical score index (CSI) recently established by our team for monitoring H5N1 infection outcomes in cats^18^. Each animal was evaluated individually, and clinical parameters such as depression, hunched posture, ruffled fur, decreased food and water intake, body weight loss, and neurological signs were recorded.^18^ Infected cats presented a CSI of 3 by day 1 post-inoculation (pi), which increased, reaching a peak average score of 18 by day 7 pi. (Fig 1B). Notably, H5N1 infection induced rapid and sustained body weight loss, averaging approximately 100 g per day beginning on day 1 pi. One infected cat (cat#11) experienced abrupt body weight loss (losing ∼12.87% of its initial body weight) and presented labored breathing by day 6 pi; this animal was therefore humanely euthanatized on day 6 pi rather than at its scheduled necropsy on day 7 pi (Figure 1C). All H5N1 infected animals developed high fever following inoculation, with body temperatures reaching approximately 41 °C on day 1 pi and remaining high (1-2 °C above baseline) through the duration of the experiment (Figure 1D). The mock-infected cats maintained or gained body weight and did not show any clinical signs throughout the experimental period (Figure 1B, 1C and 1D).

**Figure 1:**
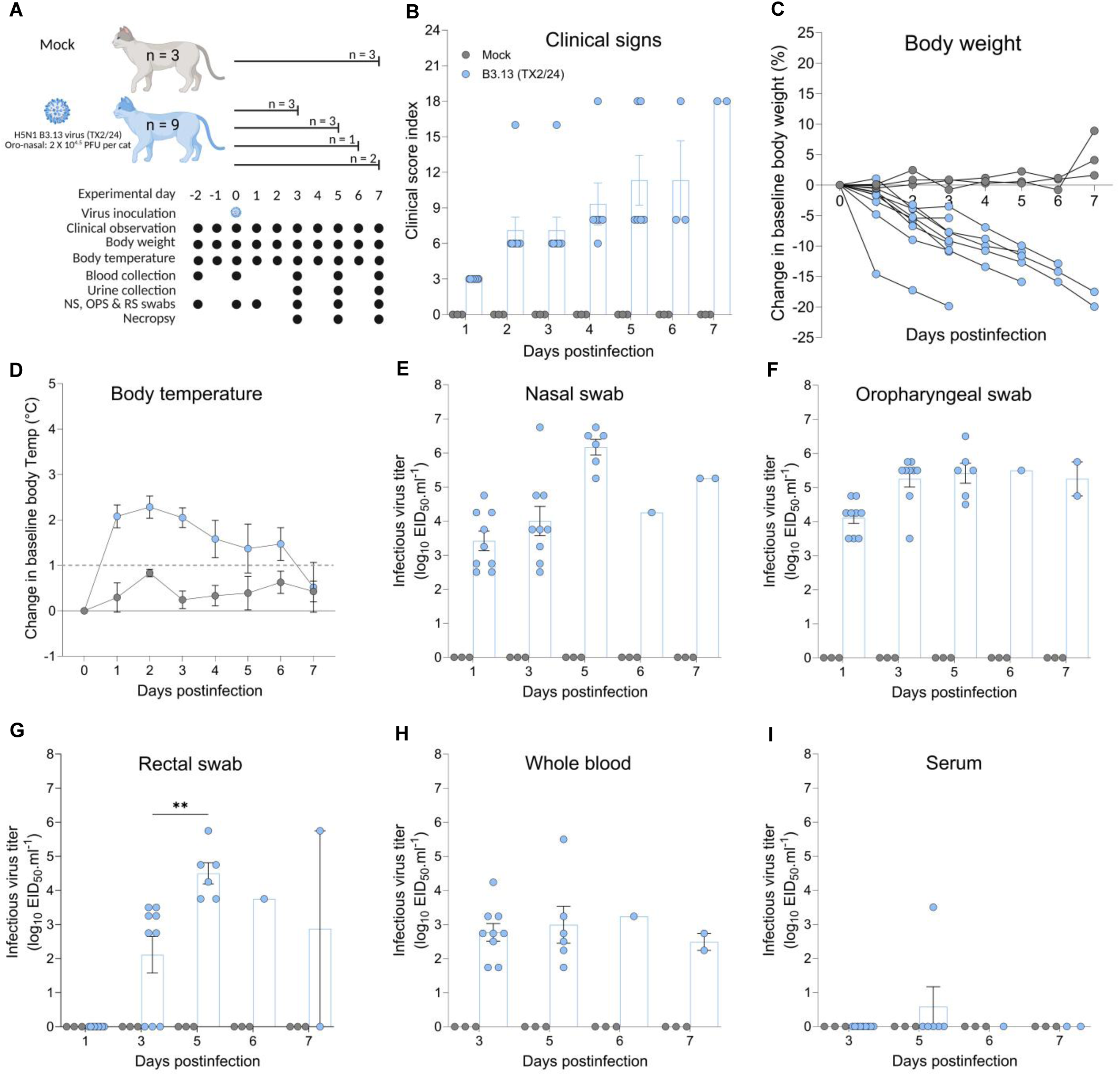
Clinical outcome of HPAI H5N1 B3.13 virus infection in cats. **A)** Experimental design of the H5N1 pathogenesis study. Twelve cats were randomly allocated into two groups of infected (n = 9) and mock (n = 3). Animals were inoculated with a virus suspension containing 10^4.5^ PFU.mL^-1^ of H5N1 B3.13 (TX2/24) through intranasal (1mL) and intraoral (1mL) routes. Animals in the mock group were inoculated with Eagle’s minimum essential medium (MEM) harvested from Cal-1 cell cultures. Three animals were euthanized on day 3 and on day 5, one on day 6, and two on day 7 (study endpoint). **B)** Cumulative daily clinical score index (CSI) recorded for mock and H5N1 B3.13 (TX2/24)-inoculated animals from 1-7 days post-infection (dpi). **C)** Changes in baseline body weight (%). **D)** Changes in baseline body temperature (°C). Infectious virus titers (log_10_EID50.ml^-1^) in **(E)** nasal **(F)** oropharyngeal **(G)** rectal swabs, **(H)** whole blood, and **(I)** serum. Each dot in the bar graphs indicates an individual animal from each group (mean with SEM, n = 3 cats/mock group, 9 cats/ H5N1 (TX2/24) infected group). Statistical significance was determined using Two-way ANOVA with column means comparison (***P < 0.0001).

### The H5N1 genotype B3.13 virus established viremia and was shed in nasal and oral secretions and in feces of infected cats

We next characterized the replication dynamics and viral shedding patterns of H5N1 B3.13 virus in cats. Nasal (NS), oral (OPS), and rectal (RS) swabs were collected on days 1, 3, 5, and 7 post-inoculation (pi) and analyzed by rRT-PCR and virus titrations in 10-day-old embryonated chicken eggs (ECEs). Analysis of virus shedding revealed high infectious viral loads in nasal and oropharyngeal secretions as early as day 1 pi (2.41-5.0 log_10_ EID_50_.mL^-1^), which peaked on day 5 pi (5.1-6.2 log_10_ EID_50_.mL^-1^) and remained high through day 7 pi (Figure 1E and 1F). Viral RNA loads in nasal and oral secretions followed similar kinetics, being first detected on day 1 (2.84 log_10_ genome copies.mL^-1^) and peaking on day 5 pi (4.2 log_10_ genome copies.mL^-1^) (Figure S1A-B). Infectious virus shedding was also detected in feces, with 6/9 animals shedding infectious H5N1 B3.13 virus on day 3 pi (3-3.9 log_10_ EID_50_.mL^-1^) with peak fecal shedding detected on day 5 pi (6/6, 4-5.8 log_10_ EID_50_.mL^-1^) (Figure 1G). Comparable kinetics were observed for viral RNA shedding (Figure S1C). Of note, viral RNA was also detected in urine from two infected cats that presented the highest clinical scores, on day 5 (cat#9) and day 6 pi (cat#11) (Figure S1D). All mock-inoculated cats remained negative throughout the study (Figure 1 and Figure S1). These results demonstrate robust replication of H5N1 B3.13 virus in cats with sustained virus shedding in oral and nasal secretions and feces of infected animals.

We also investigated the establishment of viremia following H5N1 B3.13 virus infection. High levels of infectious virus (2.8-4.3 log_10_ EID_50_.mL^-1^) (Figure 1H) and viral RNA (3.11-7.1 log_10_ genome copies.mL^-1^) (Figure S1E) were detected in whole blood of infected cats throughout the study (days 3-7 pi). Although low levels of viral RNA (1.8-4.8 log_10_ genome copies.mL^-1^) were detected in serum from 3/9 animals sampled on day 3 and 3/6 animals sampled on day 6 pi (Figure S1F), infectious virus was not recovered from serum (Figure 1I), suggesting cell-associated viremia.

### Early H5N1 B3.13 virus replication in the respiratory tract precedes systemic virus dissemination and replication in cats

We next assessed replication and tissue tropism of H5N1 B3.13 virus in infected cats. Tissues from respiratory tract (nasal turbinate, trachea and lungs), lymphoid organs (tonsils, mesenteric-, retropharyngeal-, and mediastinal lymph nodes and spleen), gastrointestinal system (stomach, duodenum, jejunum, ileum, colon, pancreas, liver), urogenital system (kidney, ovary, uterus, testes, and mammary gland), heart and nervous system (olfactory bulb, cerebrum, cerebellum and pons) were collected at necropsy performed on days 3 (n = 3), 5 (n = 3), 6 (n = 1) and 7 (n = 2) pi, and analyzed by rRT-PCR, virus titrations and *in situ* hybridization assays (Figure 2A, 2B and 2C). In respiratory tissues, high viral RNA loads were detected early during infection. High levels of viral RNA were detected in nasal turbinates between days 3-7 pi (5.78-7.5 log_10_ genome copies.g^-1^) (Figure 2A). Viral RNA loads in the trachea increased from day 3 pi (5.19 log_10_ genome copies.g^-1^) to peak levels on day 6 pi (7.52 log_10_ genome copies.g^-1^), which declined thereafter. High viral RNA loads were also observed in lung tissues (5.67-7.76 log_10_ genome copies.g^-1^) throughout the 7-day study period (Figure 2A). Moderate to high level of viral RNA loads were also detected in lymphoid tissues including tonsils, retropharyngeal-, mediastinal and mesenteric lymph nodes and spleen with peak levels observed on days 5-6 pi.

**Figure 2:**
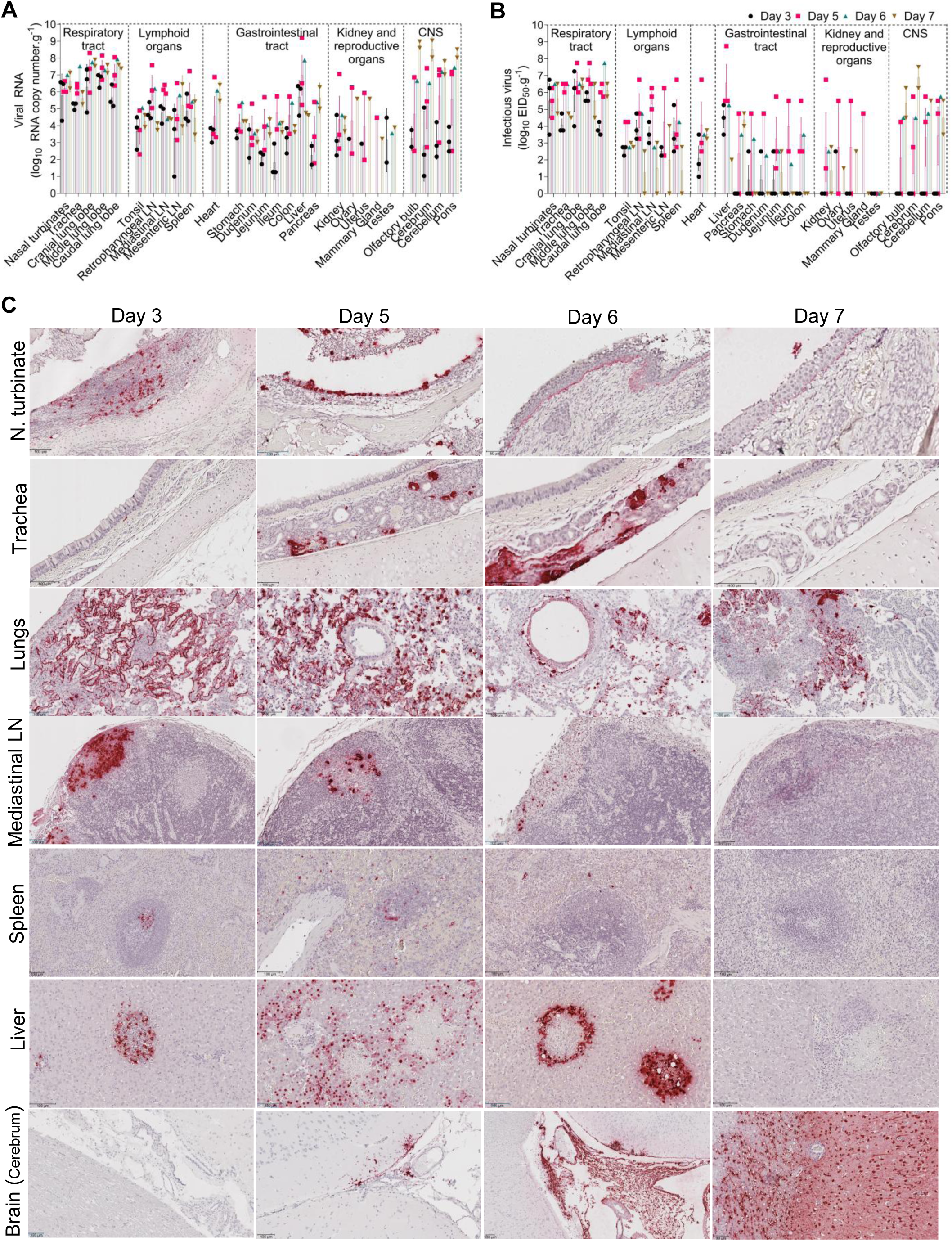
Tissue tropism, infection dynamics and systemic spread of H5N1 B3.13 virus in cats. Viral RNA **(A)** and infectious virus **(B)** loads quantified by qPCR and virus titration (EID_50_.ml^-1^) in the respiratory tract, lymphoid organs, gastrointestinal tract, kidney and reproductive organs, and brain tissues collected at necropsy days. **C)** Representative micrographs (top to bottom) of nasal turbinates, trachea, lung, mediastinal lymph node, spleen, liver, and brain from cats collected on days 3, 5, 6, and 7 post infection. In situ hybridization (ISH) of viral RNA in cells is visualized as punctate chromogenic signals (red/brown), indicating intracellular accumulation of RNA, and the cellular nuclei are counterstained with hematoxylin (blue). The ISH signals are localized to ciliated epithelial cells of the nasal mucosa, submucosal glands of the trachea, alveolar septa of the lungs, and cortical and interfollicular areas in the lymph nodes, multifocal areas of strong red labeling in the liver. In the brain, the temporal and spatial spread of viral RNA labeling is shown from microvasculature to adjacent neural tissue.

Within the gastrointestinal tract, low levels of viral RNA were detected in stomach, small and large intestine and pancreas on day 3 pi, which increased by day 5 and remained detectable thereafter (Figure 2A). In contrast, high viral RNA loads were detected in the liver as early as day 3 pi (5.71 log_10_ genome copies.g^-1^), peaking on day 5 pi (6.98 log_10_ genome copies.g^-1^) before declining on day 7 pi (Figure 2A). Lower viral RNA levels were detected in heart, kidneys and reproductive organs (ovary, uterus, mammary gland and testes) (Figure 2A).

Notably, viral RNA levels in central nervous tissues increased over the course of infection (Figure 2A). In the olfactory bulb, low viral RNA loads were detected on day 3 pi (3.25 log_10_ genome copies.g^-1^), which increased on day 5 pi (4.69 log_10_ genome copies.g^-1^) and peaked on day 7 pi (8.79 log_10_ genome copies.g^-1^). Similarly, viral RNA loads in cerebrum, cerebellum and pons increased over time as the infection progressed, with low viral RNA levels detected on day 3 pi (1.8-2.5 log_10_ genome copies.g^-1^) and peak levels observed on day 7 pi (7.43-8.54 log_10_ genome copies.g^-1^) (Figure 2A).

A similar kinetics was observed for infectious virus loads in tissues of infected cats (Figure 2B). High infectious virus titers were detected in nasal turbinates on days 3 and 5 pi (6.41-6.5 log_10_ EID_50_.g^-1^), increasing on day 6 (7.5 log_10_ EID_50_.g^-1^). In the trachea, infectious virus titers increased from day 3 pi (5.08 log_10_ EID_50_.g^-1^) to peak levels on day 5 pi (7.33 log_10_ EID_50_.g^-1^), followed by a decline on day 7 pi (5.62 log_10_ EID_50_.g^-1^). Both cranial and middle lung lobes presented very high infectious virus titers on day 3 pi (6.66-6.83), which peaked on day 5 pi (7.67-8 log_10_ EID_50_.g^-1^) and declined slightly on day 7 pi (6.5-7.5 log_10_ EID_50_.g^-1^). In contrast, the caudal lung lobes exhibited lower titers on day 3 pi (4.83 log_10_ EID_50_.g^-1^) which increased to 7.08 log_10_ EID_50_.g^-1^ and 8.75 log_10_ EID_50_.g^-1^ on day 5 and 6, respectively (Figure 2B). In lymphoid organs, initial viral replication was detected on day 3 pi (2.25-5.25 log_10_ EID_50_.g^-1^) with peak viral titers recovered from tonsil, retropharyngeal, mediastinal and mesenteric lymph nodes and the spleen on day 5 pi (2.75-6.75 log_10_ EID_50_.g^-1^), which then declined on days 6 and 7 pi (4.75 and 2.75log_10_ EID_50_.g^-1^), respectively (Figure 2B).

Within the gastrointestinal tract, low levels of infectious virus were detected on day 5 pi, which decreased thereafter. In contrast, relatively high infectious virus titers were detected in the liver on day 3 pi (5.41 log_10_ EID_50_.g^-1^), which peaked on day 5 pi (7.58 log_10_ EID_50_.g^-1^) before declining on day 7 pi (Figure 2). Low levels of infectious virus were detected in kidney and reproductive tissues throughout the experimental period (Figure 2).

Infectious virus titers in nervous tissues were first detected late in infection on day 5 pi, increasing thereafter peaking on day 7 pi (Figure 2B). Among nervous tissues, the cerebrum showed the highest infectious viral loads on day 7 pi (7.87 log_10_ EID_50_.g^-1^) followed by the cerebellum (6.5 log_10_ EID_50_.g^-1^), olfactory bulb (6.37 log_10_ EID_50_.g^-1^) and pons (5.5 log_10_ EID_50_.g^-1^).

We next localized the H5N1 B3.13 viral RNA in tissues of infected cats using *in situ* hybridization (Figure 2C). In nasal turbinates, viral RNA labeling was detected predominantly in nasal epithelium on days 3 and 5 pi. Notably, by days 6 and 7 pi, viral RNA labeling shifted primarily to subepithelial cells in nasal turbinates. In the trachea, at early time points (day 3 pi), low viral RNA labeling was detected in tracheal epithelial cells, followed by extensive viral RNA labeling within the submucosal glands on days 5 and 6 pi. On day 7 pi viral RNA labeling decreased, indicating clearance of viral RNA from the trachea. In lungs, viral RNA labeling was consistently detected along alveolar epithelial cells and in the alveolar septa on days 3 to 5 pi, followed by a marked reduction in viral labeling on day 7 pi. Viral RNA distribution in secondary lymphoid organs paralleled the systemic dissemination of the virus. Areas with strong viral RNA labeling (Figure 2C) overlapped with lymphoid necrosis in the cortex of the lymph nodes (Figure S3). In the spleen, viral RNA labeling was largely confined to germinal centers within lymphoid follicles, whereas in mediastinal lymph nodes, viral RNA labeling was detected in lymphoid follicles in the cortex and paracortex. In the liver, viral RNA labeling was detected around small multifocal necrotic foci on day 3, which expanded and coalesced by day 5 pi, with extensive RNA labeling indicative of active viral replication around the necrotic areas. Of note, cat #11, which had evidence of hepatitis at necropsy on day 6 pi had high viral RNA labeling in hepatocytes and Kupffer cell of liver (Figure 2C). Viral RNA labeling was no longer detected in the liver on day 7 pi despite persistence of necrotic foci and more extensive coagulative necrosis within the hepatic parenchyma (Figure S3).

Across the central nervous system, no viral labeling was detected on day 3 pi, while low levels of focal viral RNA labeling were detected on day 5 pi. As the infection progressed, additional viral RNA labeling was observed on day 6 with extensive viral labeling noted on day 7 pi indicating viral spread across multiple regions of the brain including cortex (parietal, occipital and temporal), hippocampus, thalamus, pons, and cerebellum. Collectively, these results revealed a temporal progression of H5N1 B3.13 virus infection, tissue tropism and replication; with the infection initiating in the respiratory tract and lymphoid organs, from where the virus disseminates hematogenously reaching multiple organs and replicating systemically before reaching the CNS and invading the brain late in the course of infection.

### Histopathological changes in tissues of cats infected with H5N1 B3.13 virus

Significant histological changes were noted in the nasal turbinates, lung, brain, liver, and lymph node, as early as day 3 pi, and were most severe in tissues collected on days 6 and 7 pi. Mild to severe neutrophil-predominant inflammation was noted in the nasal turbinates with an increase in severity of ulceration and necrosis towards days 6 and 7 pi. Changes in the lungs included mild to severe, acute interstitial pneumonia, with mixed cellularity (neutrophils, macrophages, lymphocytes, and rare plasma cells), along with necrosis, fibrin deposition, and vasculitis with fibrin thrombi. The most severe changes were noted in lung tissue collected on days 6 and 7 pi. Changes in the liver were characterized predominantly by multifocal necrosis with associated mixed inflammation and vasculitis. Coagulative necrosis was a strong feature, starting with day 3 pi, correlating with the vasculitis. Lymphoid hyperplasia and medullary histiocytosis were consistently noted in the lymph nodes, with mild lymphoid necrosis noted as early as day 3 pi.

Multifocal encephalitis, lymphoplasmacytic and histiocytic, predominantly affecting gray matter, was noted as early as day 3 pi, with additional necrosis, satellitosis, and glial nodules in tissue collected on days 6 and 7 pi. Choroid plexitis was noted in brain tissue collected on days 5 and 7 pi, likely reflective of the hematogenous route of infection, and demyelination was noted in tissue collected on day 7 pi, reflecting consequences of neuronal necrosis (Figure S3).

### Endotheliotropism of H5N1 virus leads to blood-brain barrier crossing and neuroinvasion in cats

To elucidate the pathway of H5N1 virus neuroinvasion, we examined the temporal progression of viral entry and spread within the central nervous system of infected cats. *In situ* hybridization revealed H5N1 B3.13 viral RNA labeling in brain tissue of all infected cats beginning on day 5 pi, with progressive to widespread dissemination on days 6 and 7 pi, respectively (Figure 3A). At early stages of infection (day 5 pi), viral RNA labeling was confined to endothelial cells in meningeal blood vessels, choroid plexus, cortical and periventricular blood vessels, involving the third ventricle. By day 6 pi, viral RNA labeling extended beyond the vascular endothelium into adjacent perivascular parenchyma, forming discrete perivascular foci with hybridization signals. By day 7 pi, widespread viral RNA labeling was evident in neuroparenchyma including the cortex, hippocampus, thalamus, cerebellum and pons, consistent with the extensive histopathological changes observed in these regions (Figure S2), leading to robust neuroinvasion (Figure 3).

**Figure 3:**
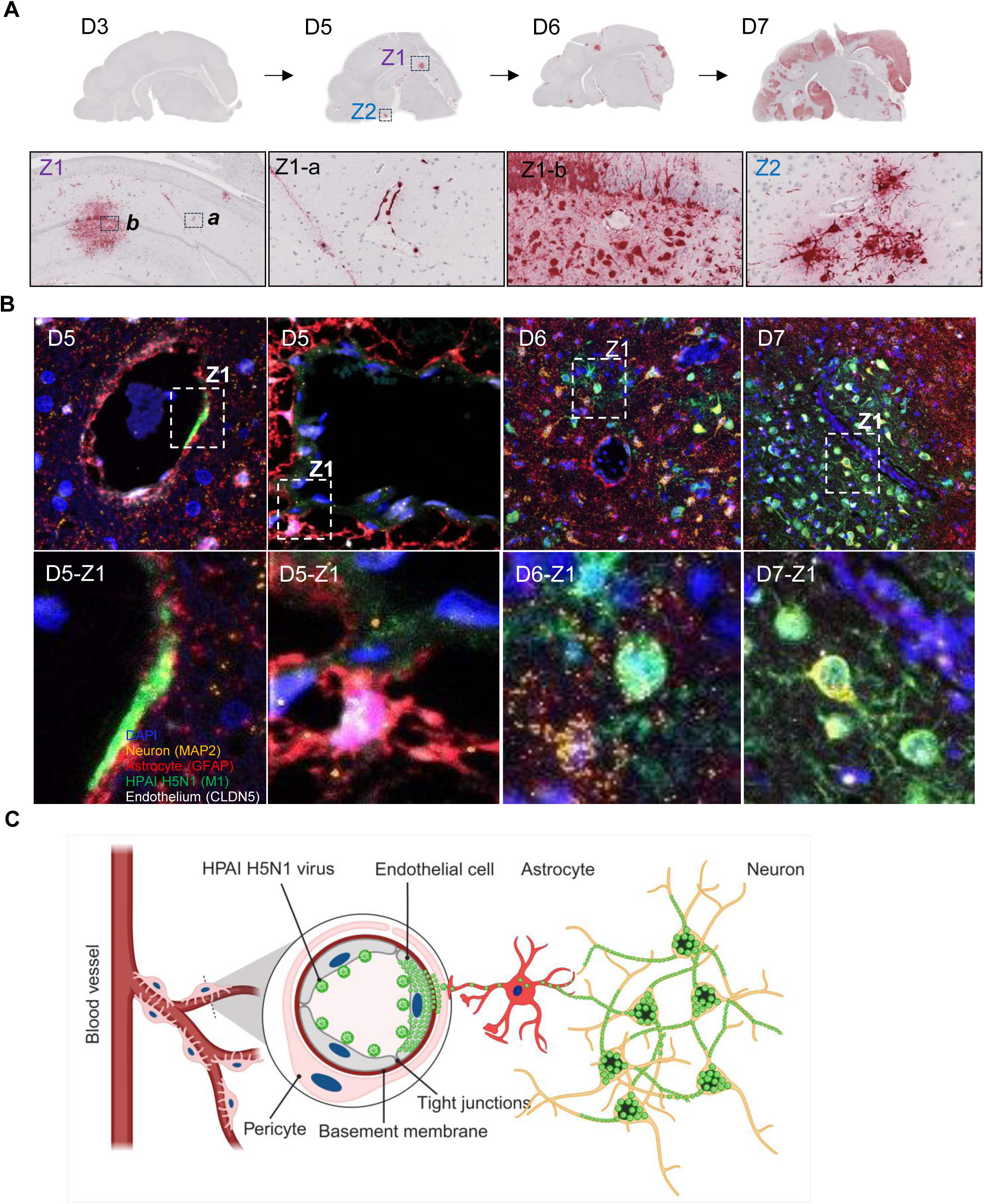
Neuroinvasion of H5N1 B3.13 virus through blood-brain-barrier crossing. **A)** Spatial and temporal distribution of H5N1 B3.13 virus RNA in the brain, demonstrating hematogenous entry and subsequent focal spread. Brain sections collected on days 3, 5, 6, and 7 pos infection reveal viral RNA localized to endothelial cells and perivascular regions of both small and large vessels, indicating hematogenous neuroinvasion. Higher-magnification areas (Z labeled) show viral RNA labeling within vascular walls and adjacent parenchyma, consistent with transendothelial spread. **B)** Brain tissues were processed by fluorescent in situ hybridization (FISH) using a viral probe targeting the M1 gene (green). Cell-type–specific markers included CLDN5 for endothelial cells (white), GFAP for astrocytes (red), and MAP2 for neurons (orange). Colocalization of viral RNA with endothelial cells, astrocytes, and neurons is indicated by the overlap of green with white, red, and orange colors, respectively, and is shown in higher magnification (Z labeled areas). **C)** Schematic diagram depicting neuroinvasion by H5N1 virus through the blood-brain barrier. The virus initially infects endothelial cells of the brain microvasculature, spreading to adjacent astrocytes which ensheathe the microvasculature, and finally spreading to neurons

Based on the sustained viremia detected in infected animals and early endothelial localization of viral RNA, we hypothesized that H5N1 virus reaches the brain via hematogenous dissemination. To test this hypothesis, we further characterized the cellular tropism and temporal sequence of neuroinvasion using a 4-plexed fluorescent *in situ* hybridization (FISH) approach incorporating a virus-specific probe and markers for endothelial cells (Claudin-5; CLDN5), astrocytes (Glial fibrillary acidic protein; GFAP) and neurons (Microtubule-associated protein 2; MAP2) applied to brain tissues collected at necropsy on days 3, 5, 6 and 7 pi (Figure 3B, Figure S4). No viral RNA was detected in brain tissues on day 3 pi. The earliest detection of hybridization signal occurred on day 5 pi (Figure 3B), when viral RNA localized predominantly within vascular endothelial cells (Figure 3B). Given the close anatomical association between endothelial cells and astrocytes, viral RNA labeling was frequently detected in adjacent perivascular astrocytes, as indicated by colocalization of viral RNA (green) with GFAP-positive astrocytes (red) (Figure 3B, D5-Z1). In contrast, astrocytes distal to blood vessels were infrequently infected (Figure 3B). As infection progressed, the virus spread extensively into the cortex, with neurons representing the most abundantly infected cell population (Figure 3b, D7-Z1). These results demonstrate that H5N1 virus crosses the blood-brain barrier to invade and replicate in neurons(Figure 3C).

### The H5N1 B3.13 virus is more pathogenic and transmissible than the D1.1 virus in cats

Following the introduction of Eurasian H5N1 clade 2.3.4.4b viruses into North America in late 2021, multiple reassortant genotypes comprising Eurasian and North American lineage gene segments rapidly emerged and became established in wild bird reservoirs across the United States^3^. The B3.13 genotype emerged in late 2023^10^ through reassortment of an ancestral Eurasian genotype A1 virus. This genotype retained four Eurasian lineage gene segments (PA, HA, NA and M) while acquiring four gene segments-PB2 [am2.2], PB1 [am4], NP [am8] and NS [am1.1]-from North American low pathogenic avian influenza (LPAI) viruses (Figure 4A). The B3.13 genotype subsequently became predominant in dairy cattle and was associated with extensive spillovers into mammals, including cats and humans. ^5,10^ In contrast, the D1.1 genotype emerged in wild birds in the fall of 2024 as a reassortant derived from the fully Eurasian A3 genotype. The D1.1 genotype viruses retained four Eurasian gene segments (PB1, HA, M, NS) while incorporating four LPAI North American lineage segments-PB2 [am24], PA [am4], NP [am13] and a novel NA [am4]) segment (Figure 4A). Notably, acquisition of the North American N1 segment represented a major antigenic shift, replacing the previously fixed Eurasian N1. We analyzed publicly available nucleotide sequences (GISAID)^10^ from B3.13 and D1.1 viruses recovered in the US between 2023 and 2025 and assessed the association of these two major genotypes across different animal species, including avian (chickens) and mammalian hosts (including cattle, cats and humans). While viruses from both genotypes have been detected in all four species, H5N1 B3.13 viruses have been more frequently detected in cattle, cats, and humans when compared to the D1.1 viruses. Interestingly, the frequency of detection of B3.13 viruses in chickens was lower than for D1.1 viruses (Figure 4B).

**Figure 4:**
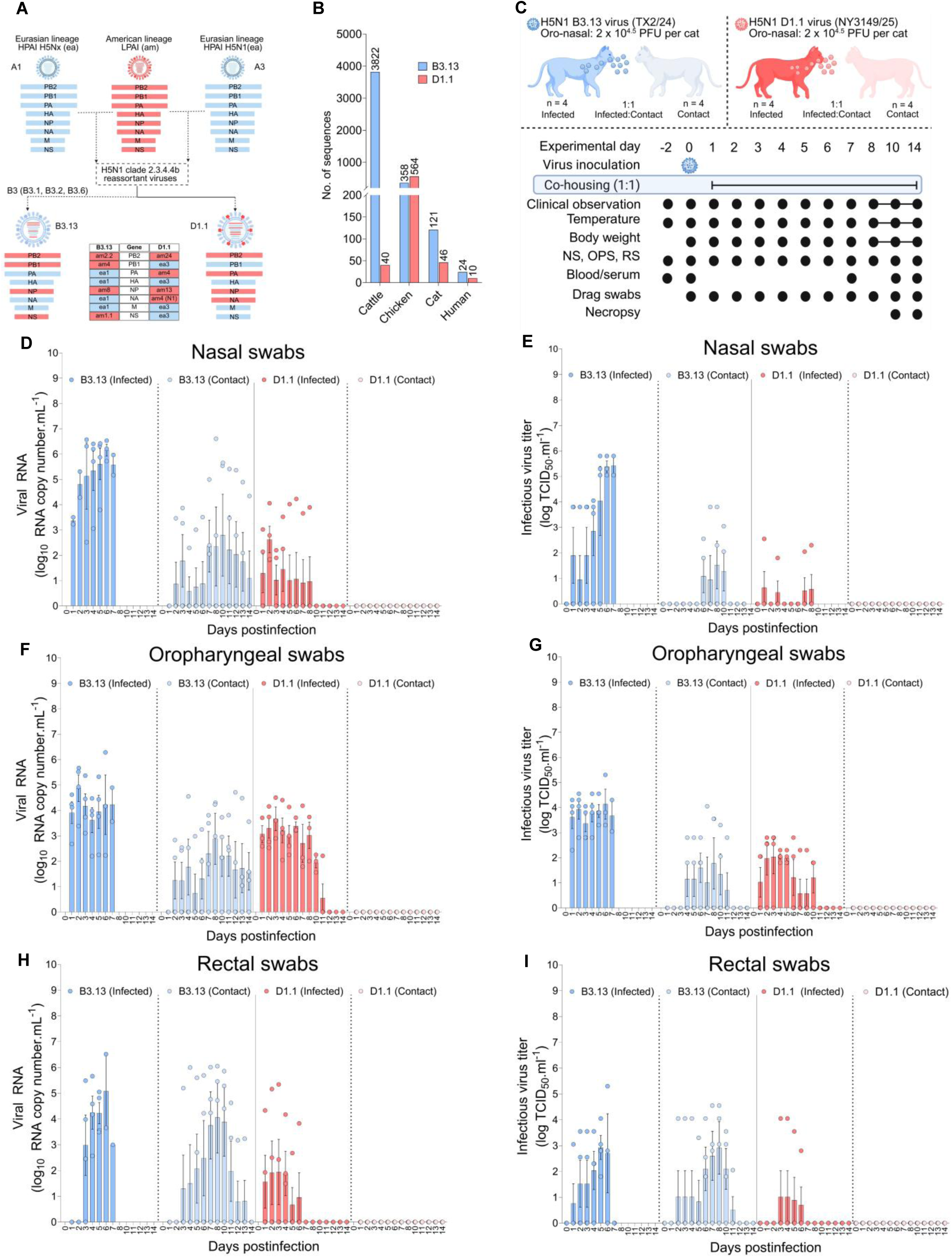
Transmission efficiency of H5N1 B3.13 and D1.1 genotypes in cats. **A)** A schematic diagram depicting origin of H5N1 B3.13 and D1.1 genotypes following reassortment of parental Eurasian A1 and A3 genotypes with American lineage low pathogenic avian influenza (LPAI) viruses, respectively. Blue bars represent Eurasian and orange bars depict American lineage gene segments. **B)** Detection frequency and distribution of H5N1 B3.13 and D1.1 genotypes in cattle, chicken, cat and human hosts. **C)** Direct contact transmission study design. Infected cats were inoculated with 10^4.5^ PFU.ml^-1^ of H5N1 B3.13 (TX2/24) or D1.1 (NY3149/25) through the intranasal (1mL) and intraoral (1mL) routes. At 24h post-inoculation infected cats were moved to an enclosure with the naïve contact animals (1:1 ratio). Infected and contact animals were co-housed for the remaining of the study to assess direct-contact transmission. Nasal, oropharyngeal, and rectal swabs were collected daily from both infected and contact animals. Clinical signs, body weight and temperature were monitored for the study duration. **c,** Viral RNA (quantified by qPCR) and infectious virus titers (TCID_50_.ml^-1^) in nasal **(D, E)**, oropharyngeal **(F, G)**, and rectal swabs **(H, I)**. H5N1 B3.13 virus exhibited higher viral RNA and infectious virus shedding and greater transmission to contacts compared with genotype D1.1 virus. qPCR quantification of H5N1 RNA in nasal, oropharyngeal, and rectal swabs. B3.13-infected index cats shed higher levels of viral RNA compared to D1.1. Contact cats exposed to B3.13 also showed viral RNA detection but not contact cats from D1.1. Each dot in the bar graphs indicates an individual animal from each group (B3.13: Infected n = 4, Contact n = 4; D1.1: Infected n = 4, Contact n = 4).

Next, we compared the pathogenicity, replication properties, and transmission efficiency of B3.13 and D1.1 viruses in cats using a direct contact transmission (DCT) model. For this, 8 cats were inoculated oronasally with 2 x 10^4.5^ PFU of either B3.13 (TX2/24) (n = 4) or D1.1 (NY3149/25) (n = 4) virus and 24 h later co-housed with naïve contact cats (1:1 ratio) for the duration of the experimental period (Figure 4C). Cats infected with either virus developed moderate-to-severe disease, characterized by elevated clinical score index (CSI), fever and rapid weight loss (Figure S5A-D). However, disease progression was faster, and the severity was greater in animals inoculated with the B3.13 (TX2/24) virus, with all infected cats reaching humane endpoints by day 7 pi (100% mortality) (Figure S5E). In contrast, infection with the D1.1 (NY3149/25) virus resulted in delayed disease progression, with 3 of 4 cats euthanized by day 12 pi (75% mortality) (Figure S5E).

Notably, transmission outcomes differed markedly between the two virus genotypes. Two of four contact cats exposed to B3.13 (TX2/24)-infected animals developed clinical disease as evidenced by increased CSI scores, loss in body weight and fever, whereas all contact cats exposed to D1.1 (NY3149/25)-infected animals remained clinically normal and continued to gain weight throughout the 14-day study period. We also quantified infectious virus and viral RNA shedding in nasal (NS, nasal swabs), and oral (OPS, oropharyngeal swabs) secretions and feces (RS, rectal swabs) of infected and contact cats using virus titrations and rRT-PCR. Across all sample types, cats infected with B3.13 (TX2/24) virus exhibited consistently higher viral RNA and infectious virus loads than cats infected with D1.1 (NY3149/25) throughout the experimental period (Figure 4D-I). In nasal secretions, viral RNA levels in B3.13 (TX2/24)-infected cats increased rapidly from day 1 pi and peaked on day 6 pi (6.15 log_10_ genome copies.mL^-1^), whereas viral RNA levels in D1.1 (NY3149/25)-infected cats were lower in magnitude and transient. Three of four (3/4) contact cats exposed to B3.13 (TX2/24)-infected animals shed viral RNA at lower but detectable levels over the 14-day experimental period, while no nasal shedding was detected in contact cats exposed to D1.1 (NY3149/25)-infected animals. Infectious virus titers in nasal secretions mirrored RNA kinetics: B3.13 (TX2/24)-infected and contact cats shed infectious viruses as early as days 1 and 6 pi, respectively, whereas infectious virus in nasal secretions were detected only sporadically in one D1.1 (NY3149/25)-infected cat and it was not detected in D1.1-contact animals (Figure 4E). Oral shedding of B3.13 (TX2/24) virus was high, peaking on day 6 pi (4.2 log_10_ genome copies.mL^-1^), with detectable shedding in 3/4 contact cats (Figure 4F). In contrast, oral shedding of D1.1 (NY3149/25) virus was lower (3.38 log_10_ genome copies.mL^-1^) on day 6 pi, and was not detected in contact animals. High infectious virus titers were detected in oral secretions from all B3.13 (TX2/24)-infected cats (3/4) and intermittently in 3 out of 4 contact cats, whereas infectious viral titers in D1.1 (NY3149/25)-infected cats were lower (1.8-2.3 log_10_ TCID_50_.mL^-^) on day 5 and were undetectable in contact animals (Figure 4G). Fecal shedding followed a similar pattern, where B3.13 (TX2/24)-infected cats shed high viral RNA loads in feces which peaked on day 6 pi, with substantial viral RNA shedding in contact cats, whereas D1.1 (NY3149/25)-infected cats exhibited lower and transient RNA shedding in feces, and contact cats remained negative throughout the study (Figure 4H). Infectious viruses were detected in fecal samples from all B3.13 (TX2/24)-infected and 3 out of 4 contact cats, while only 1 out of 4 D1.1 (NY3149/25)-infected cats shed infectious viruses in feces and none of the contact animals shed infectious virus in feces (Figure 4I). Efficient transmission of the B3.13 (TX2/24) virus from infected to contact animals was confirmed by virus neutralization assays demonstrating seroconversion of 3/4 contact animals. In contrast, none of the contact cats in the D1.1 (NY3149/25) group seroconverted, indicating a lack of virus transmission (Figure S5F).

Consistent with the shedding patterns in respiratory secretions and feces from infected and contact animals, viral RNA was detected throughout the experiment in environmental swab-(enclosure-, feeder- and waterer swabs) and water samples from enclosures housing the B3.13 (TX2/24)-infected:contact cats. No viral RNA was found in environmental samples from the D1.1 (NY3149/25)-infected:contact animals. Infectious virus was intermittently recovered from enclosure swabs from the B3.13 (TX2/24)-group between days 2-13 pi, from feeder and waterer swabs between days 7-8 pi and from water samples collected on day 7 pi, while no infectious virus was isolated from environmental samples from the D1.1 (NY3149/25) group (Figure S7). Collectively, these results demonstrate a higher pathogenicity and transmissibility of H5N1 B3.13 (TX2/24) genotype virus when compared to D1.1 (NY3149/25) virus in cats.

### Replication kinetics of H5N1 B3.13 and D1.1 viruses *in vitro*

Given the differences in virus replication and shedding between H5N1 B3.13 and D1.1 viruses observed *in vivo*, we next compared their replication and virus release kinetics in mammalian (feline, bovine and human) and avian (chicken) cell cultures *in vitro*. For this, multi-step growth curves (MOI = 0.1) were performed in feline lung (FelLung), bovine uterine epithelial cells (Cal-1), human alveolar epithelial cells (A549), and chicken fibroblast cells (DF-1) and virus titers determined at 4, 8, 12, 24, 48 and 72 hpi in cell supernatant and the cell pellet to quantify cell free and cell associated viruses, respectively. In mammalian cells, the B3.13 (TX2/24) virus replicated more efficiently than the D1.1 (NY3149/25) virus, as reflected by consistently higher infectious virus titers (0.5-2 log_10_ TCID_50_.mL^-1^) recovered from both cell culture supernatants and cell-associated fractions (Figure 5A-H). In contrast, both viruses exhibited comparable replication kinetics with only slight differences (∼ 0.5-1 log_10_ TCID_50_.mL^-1^) observed in the cell-associated fraction in chicken-derived DF-1 cells, indicating similar fitness in avian cells. These results suggest a host-specific replication advantage of the H5N1 B3.13 virus in mammalian cells.

**Figure 5:**
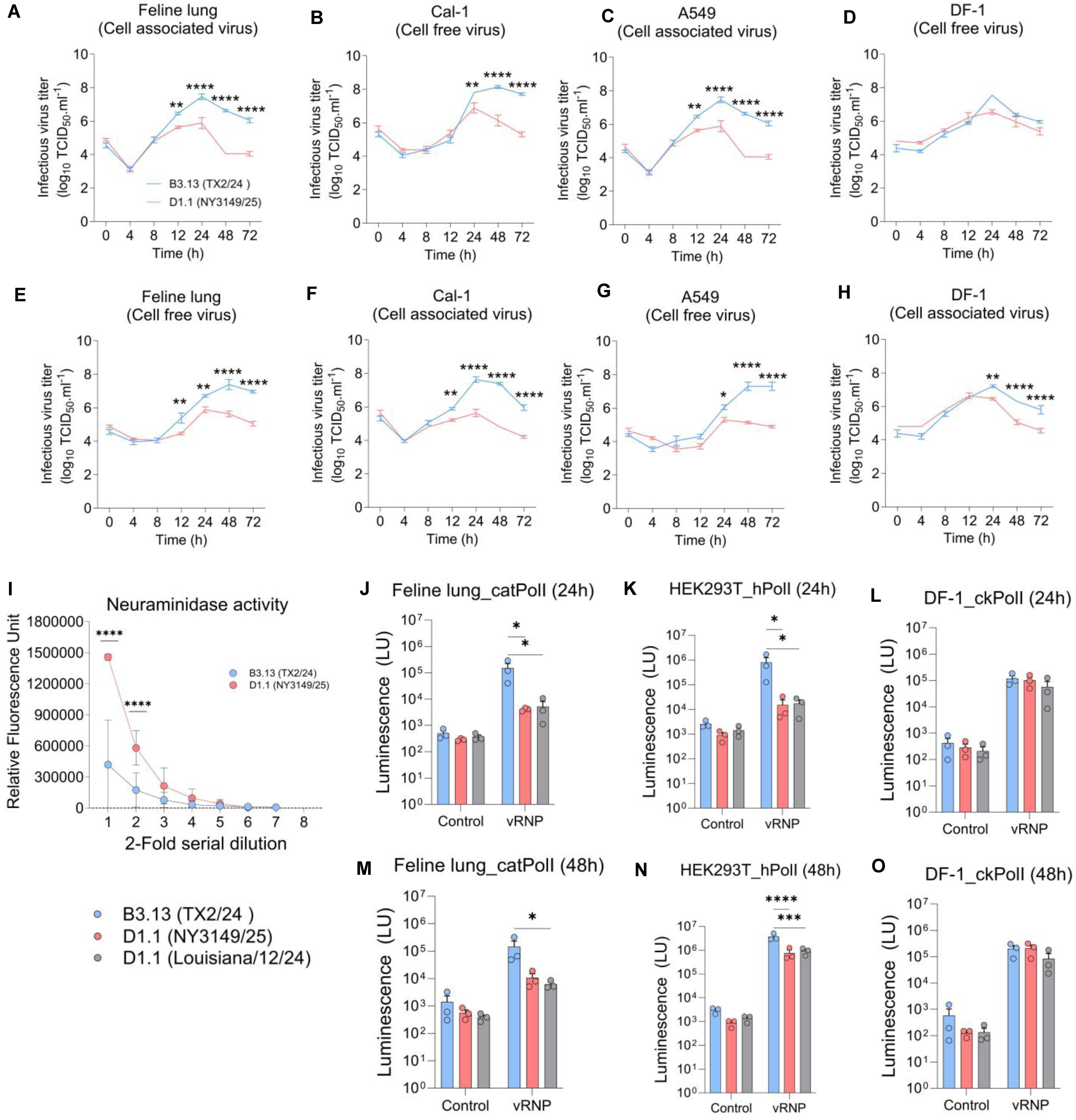
Replication efficiency of H5N1 B3.13 and D1.1 genotypes. Replication kinetics of H5N1 B3.13 and D1.1 viruses were evaluated in feline lung cells **(A, E),** bovine uterine epithelial Cal-1 cells **(B, F),** human lung epithelial A549 cells **(C, G),** and chicken fibroblast DF-1 cells **(D, H).** Monolayers of cells were infected in triplicate at a multiplicity of infection (MOI) of 0.1 and incubated at 37 °C with 5% CO₂ for 1 h. Cell culture supernatants were collected at 0, 4, 8, 12, 24, 48, and 72 h post-inoculation to quantify cell-free virus, while corresponding cell lysates were harvested to assess cell-associated virus. Viral titers were determined by calculating the 50% tissue culture infectious dose (TCID₅₀ ml⁻¹) using the Spearman–Kärber method and are expressed as log₁₀ (TCID₅₀ mL⁻¹). **(I)** Neuraminidase (NA) activity of H5N1 B3.13 and D1.1 viruses was quantified using a fluorometric NA activity assay (NA-Fluor, Applied Biosystems). Virus stocks were diluted (2-fold dilutions) to a baseline concentration of 5 × 10⁷ PFU mL⁻¹ . Assays were performed in triplicate. Background-subtracted relative fluorescence units (RFU) were plotted against virus dilution, and the dilution yielding RFU values within the linear range of the 4-methylumbelliferone (4-MU) standard curve were selected for normalization across the H5N1 B3.13 and D1.1 viruses. Polymerase activity of H5N1 B3.13 (TX2/24) and D1.1 (NY3149/25 and LA/12/24) genotypes was determined in feline lung epithelial cell **(J, M)**, HEK293T cells **(K, N),** and DF-1 cells (**L, O)**. Cells were co-transfected with pCAGGS plasmids encoding the genotype-specific viral ribonucleoprotein (RNP) complex components PB2, PB1, PA, and NP, and with the pUC57-NP (Non-coding region [NCR])-NLuc minigenome reporter plasmids. Control cells were transfected with the vRNP complex proteins and the pUC57-NP minigenome plasmid minus the PB1 encoding plasmid. Nanoluciferase activity was analyzed by luciferase reporter assay. Data represent the mean ± SEM from three independent experiments.

### The neuraminidase (NA) activity of H5N1 B3.13 virus is lower than that of the D1.1 virus

One of the major differences between H5N1 genotypes B3.13 and D1.1 viruses lies in the neuraminidase (NA) gene segment. While B3.13 viruses retained the Eurasian N1 segment that emerged in Europe in 2020-2021 in the ancestral H5N1 A1 genotype viruses^3,19^, D1.1 genotype viruses acquired a North American LPAI-derived N1 segment through reassortment. To assess the functional impact of this reassortment and investigate whether the neuraminidase activity could help explain differences in the transmissibility of these viruses, we compared the B3.13 (TX2/24) and the D1.1 (NY1349/25) N1 protein sequences (Supplementary Table 2) and their neuraminidase activities (Fig 5I). The two N1 proteins differed by 29 amino acid residues distributed across all major functional domains, including 12 substitutions in the transmembrane and stalk region (positions 1-84, N1 numbering), which influence NA stability, and 17 substitutions in the globular head domain (positions 85+, N1 numbering), which contain the conserved catalytic site (R119, R292 and R371). To determine whether these sequence differences are translated into functional divergence, we measured the neuraminidase activity of the B3.13 (TX2/24) and D1.1 (NY3149/25) NAs using the fluorogenic NA substrate 4-MUNANA. The B3.13 (TX2/24) virus NA consistently exhibited lower neuraminidase activity than the D1.1 NY1349/25 virus NA, as indicated by increased relative fluorescence units (RFU) (Figure 5I). These findings suggest that NA is not the major driver underlying the differences in pathogenicity and transmissibility between B3.13 (TX2/24) and D1.1 (NY3149/25) viruses observed in our study.

### Higher polymerase activity of H5N1 genotype B3.13 than D1.1 viruses

The emergence of H5N1 B3.13 and D1.1 genotypes involved extensive reassortment within the viral ribonucleoprotein (vRNP) complex, comprising the polymerase subunits (PB2, PB1, PA) and NP, between Eurasian H5N1 and North American LPAI viruses.^3^ Comparative analysis of the amino acid sequences revealed substantial divergence between B3.13 (TX2/24) and D1.1 (NY3149/25), including differences in PB2 (9 substitutions), PB1 (14), PA (18), and NP (3) proteins (Supplementary Table 2). Although neither virus encodes the canonical mammalian adaptive PB2 E627K mutation, the bovine-derived B3.13 (TX2/24) virus carries the PB2 M631L substitution. To assess the functional consequences of these differences, we compared their polymerase activities using minigenome replicon assays. Polymerase complexes derived from B3.13 (TX2/24) and D1.1 (NY3149/25), or a human-derived D1.1 virus (A/human/Louisiana/12/2024; Louisiana/12), were evaluated in feline lung (FelLung), human (HEK293T), and avian (DF-1) cells. The B3.13 (TX2/24) polymerase complex exhibited significantly higher activity than both D1.1 (NY1349/25 and Louisiana/12) polymerase complexes in feline as well as in human cells, as indicated by increased luciferase activity (p ≤ 0.05) (Figure 5J-O). In contrast, polymerase activities of B3.13 (TX2/24) and D1.1 (NY3149/25 and Louisiana/12) viruses were comparable in chicken-derived DF-1 cells. Minimal background signals were observed in negative control cells lacking PB1. These results indicate that the higher efficiency of the B3.13 (TX2/24) vRNP complex could contribute to its enhanced replication ability in feline cells observed both *in vitro* and *in vivo* in the present study.

## Discussion

The panzootic H5N1 clade 2.3.4.4b viruses have undergone remarkable genetic diversification in recent years, with reassortment with LPAI viruses in reservoir waterfowl and shorebirds being the main driver of virus evolution.^3,20^ This resulted in major host range expansion (with over 500 species affected) – leading to spillover to mammalian species in close contact with humans (dairy cows and cats). Additionally, diverse clinical manifestations varying from respiratory, neurologic, and systemic disease to mastitis have been reported.^5,9,17^ Domestic cats are among the most severely affected species with animals often presenting severe respiratory and/or neurological diseases, usually succumbing to infection. In this study, we investigated the infection dynamics, tissue tropism, and the pathway of neuroinvasion of H5N1 virus infection in cats. Our results show that following oronasal exposure, H5N1 virus replicates in the upper respiratory tract epithelium, establishing rapid and sustained cell-associated viremia^21^, which is followed by dissemination to and replication in lungs and trachea, lymphoid organs (tonsil, lymph nodes, and spleen), liver, and the gastrointestinal tract, causing systemic infection, leading to multisystemic cell- and tissue damage. Replication of H5N1 virus in this broad array of organs coincided with peak viremia levels^21^, suggesting the occurrence of secondary viremia following virus replication in visceral and lymphoid organs and subsequent viral spread to the central nervous system (CNS) later in infection (Figure 6).

**Figure 6:**
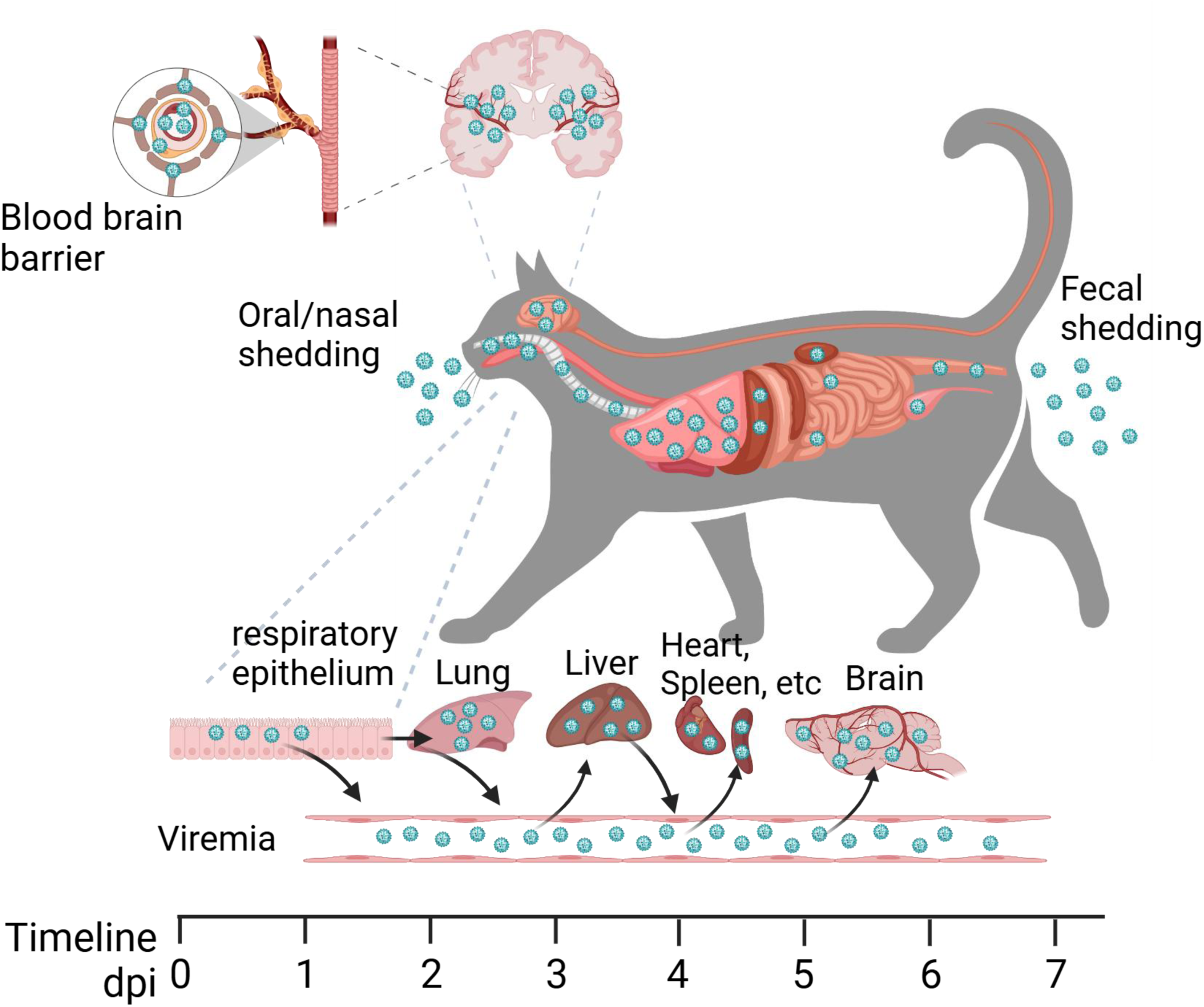
H5N1 virus infection dynamics and pathogenesis in cats. The schematic illustrates the temporal and systemic progression of H5N1 infection following oronasal exposure in cats. The virus replicates initially in the epithelium of the upper respiratory tract (days 1–2 postinfection), spreads to the lungs, and establishes early viremia. The circulating virus disseminates to other sites of virus replication including, liver, spleen, and heart. Virus shedding in respiratory and oral secretions initiates on day 1 and is detected until day 7 pi, while virus shedding in feces is observed between days 3-5 pi. By days 5–7, the virus reaches the brain. Neuroinvasion occurs via crossing of the blood-brain barrier of the brain neurovasculature, followed by infection of glial cells (astrocytes and neurons). Central nervous system infection results in perivascular multifocal to diffuse spread of the virus.

The broad tissue tropism of H5N1 virus observed in felines is a hallmark of the virus infections in mammalian hosts, such as foxes, raccoons, and mink.^2,22,23^ However, this contrasts strikingly with the virus tropism observed in dairy cows, where viral replication is largely restricted to the mammary gland, resulting in localized infection and severe mastitis.^5,24,25^ One key difference in H5N1 pathogenesis that may explain the disseminated systemic replication seen in cats versus the localized infection in dairy cows is the establishment of robust and sustained viremia in cats, as demonstrated here and in our previous study^21^, compared to the absence of viremia or only transient, low-level viremia in dairy cows.^26–29^ The differential distribution of α2,3 and α2,6 sialic acid receptors may also contribute to these distinct patterns of virus tropism across different susceptible species.

A unique feature of HPAI H5Nx viruses, and of the currently circulating H5N1 clade 2.3.4.4b viruses, is their ability to cause severe neurological disease in avian and mammalian hosts. In cats, a marked neurotropism has been described in several cases of natural H5N1 infections.^28,30,31^ This has been associated with abundant expression of α2,3 and α2,6 sialic acid receptors in the cat brain, including the cortex, hippocampus, cerebellum and brain stem.^31^ Importantly, the widespread distribution of viral receptors in the brain correlates with the sites of viral replication observed here and described in cases of natural infection.^28,30,31^ Despite this, certain aspects of the neuropathogenesis – including the routes/pathways of neuroinvasion – of H5N1 viruses in mammalian hosts remain elusive. Studies in mice and ferret models suggest that H5N1 viruses can use either the olfactory pathway through sensory neurons that innervate the nasal mucosa, or the respiratory pathway through the trigeminal, facial and glossopharyngeal nerves in the upper respiratory tract or yet through the vagus nerve in the lower respiratory tract to enter the CNS.^32,33^ Here we showed that in cats, B3.13 (Figure S2) and D1.1 (Figure S6) viruses invade the CNS via the hematogenous pathway after crossing the blood-brain barrier. Following a viremic phase, the virus reaches the brain microvasculature, where it initially infects endothelial cells before spreading to astrocytes. These astrocytes ensheathe the vascular tube and connect blood vessels to neuronal circuits in the brain parenchyma, leading to viral spread, infection, and replication in neurons. The multifocal distribution of H5N1 virus replication across various anatomic regions of the CNS demonstrates that this entry pathway enables efficient neuroinvasion, resulting in multifocal meningoencephalitis. These observations indicate that the hematogenous pathway likely plays a more important role in H5N1 viral neuropathogenesis than previously recognized.^34^ This insight has important implications for developing targeted therapeutic interventions against H5N1 infection in mammals, as effective treatment of viral encephalitis may require the use of drugs that cross the blood-brain barrier to contain viral replication in the brain.

The primary concern regarding the rapid genetic diversification observed in H5N1 viruses is the potential emergence of variants with enhanced transmissibility or pathogenicity. Epidemiological evidence indicates that both H5N1 genotypes B3.13 and D1.1 exhibit improved fitness when compared to their ancestral strains. The B3.13 genotype successfully spilled over into dairy cattle, establishing efficient transmission chains within this novel host species and subsequently spreading to other mammalian- (including humans and cats) and avian hosts. Similarly, the D1.1 genotype emerged in wild birds, causing the largest infection wave since the introduction of H5N1 in North America. This genotype also crossed the species barrier, jumping from wild birds to dairy cows in three independent spillover events across Nevada, Arizona, and Wisconsin.^10,35^

When comparing the frequency of complete genome sequences of H5N1 B3.13 and D1.1 genotypes identified across mammalian and avian (chickens) species, we observed a higher detection rate of B3.13 genotype viruses in mammalian hosts including cattle, cats and humans, while D1.1 sequences were more frequently detected in chickens. Our experimental results from *in vivo* studies in cats and *in vitro* analyses across mammalian and avian cell lines demonstrate enhanced fitness of the H5N1 B3.13 (TX2/24) virus in mammalian hosts, as evidenced by increased replication efficiency. Conversely, both B3.13 (TX2/24) and D1.1 (NY3149/25) viruses exhibited comparable replication rates in chicken cells. However, it is important to note that the D1.1 (NY3149/25) virus utilized is an avian- derived isolate. Whether other D1.1 isolates – particularly those recovered from mammalian hosts – present the same properties remains to be determined.

The enhanced transmissibility of the B3.13 (TX2/24) virus observed in our direct contact transmission (DTC) studies in cats corroborates the virus shedding and replication levels detected in this group, compared to D1.1 (NY3149/25)-infected cats. Notably, virus replication and shedding levels in nasal and oral secretions, as well as in feces of infected and contact cats, correlated with viral loads detected in environmental samples collected from the animal enclosures, feeder, and water. While environmental contamination was readily detected in enclosures housing B3.13 (TX2/24)-infected cats, no virus was detected in enclosures from D1.1 (NY3149/25)-infected animals. These findings suggest that the B3.13 (TX2/24) virus transmission could occur through multiple routes, including direct contact between infected and contact animals via respiratory or oral aerosols shed by infected animals, or indirect contact with contaminated environmental fomites (feed, water or surfaces). Importantly, enhanced direct contact transmission of B3.13 compared to D1.1 viruses (human isolates) has also been observed in a ferret model^36^, highlighting the higher transmission risk of the B3.13 genotype in mammalian hosts.

Transmission of H5N1 virus is multifactorial and can be influenced by the virus’ affinity for host cell receptors, replication levels, and efficiency of progeny virus egress. These functions are coordinated by several key viral proteins: the HA and NA affect viral interactions with host cell receptors, while the polymerase subunits (PB2, PB1, and PA) are responsible for viral replication. Sequence comparison between B3.13 and D1.1 viruses revealed that while both genotypes retained the same HA gene segment, important differences in other gene segments were observed. For example, the B3.13 viruses retained the NA gene from the Eurasian H5N1 lineage, while the D1.1 genotype viruses acquired the NA gene from North American LPAI lineage viruses. Given the differences in NA sequences between B3.13 and D1.1 viruses, we analyzed and compared the enzymatic activity of each genotype’s neuraminidase. These experiments revealed a higher NA activity of the D1.1 protein, compared to the B3.13 virus, indicating that the lower viral loads detected in respiratory secretions and feces from D1.1-infected cats cannot be attributed to impaired NA activity or compromised virus release from infected cells.

Major sequence variations were also observed in the polymerase complex proteins (PB2, PB1, PA) of the B3.13 and D1.1 viruses. Thus, we evaluated and compared the polymerase activity of each genotype using replicon systems, where replication of a minigenome cassette encoding a nano-luciferase reporter was assessed in a panel of cells derived from mammalian (cats and humans) and avian (chickens) hosts. These studies revealed a higher polymerase activity of the B3.13 (TX2/24) vRNP complex in feline and human-derived cells compared to the vRNP complexes of two D1.1 viruses (NY3149/25-avian origin, and Louisiana/12/24-human origin), while vRNP complexes of both genotypes presented similar activity in chicken cells. These findings align with the replication kinetics of the authentic B3.13 (TX2/24) and D1.1 (NY3149/25) viruses observed in cats *in vivo* and in mammalian and avian host cell lines *in vitro*. Collectively, our results suggest that differences in the polymerase activities between B3.13 (TX2/24) and D1.1 (NY3149/25) viruses may contribute to their distinct abilities to replicate and transmit in the cat host. Further investigations are needed to determine whether both viruses replicate to similar levels in avian hosts in vivo. Our study revealed critical aspects of H5N1 virus pathogenesis in a naturally susceptible animal species, emphasizing the risk posed by B3.13 viruses currently circulating in dairy cattle in the U.S. and frequently transmitted to cats.

## Methods

### Biosafety and ethics statement

All work related to handling, propagation, and characterization of highly pathogenic avian influenza (HPAI) H5N1 viruses was performed under strict biosafety measures at the Animal Health Diagnostic Center BSL3 research suite at the College of Veterinary Medicine, Cornell University. All animals were handled in accordance with the Animal Welfare Act (AWA). The study procedures were reviewed and approved by the Institutional Animal Care and Use Committee (IACUC) at Cornell University (Protocol no. 2024-0094).

### Cells and Virus

Bovine uterine epithelial cells (Cal-1; developed in house in virology laboratory at Animal Health Diagnostic Center [AHDC]), human lung carcinoma cells (A549), chicken embryo fibroblast cells (DF-1), Feline lung cells (FelLung) and Madin-Darby canine kidney (MDCK) cells were propagated in minimum essential media (MEM, Corning. In, Corning, NY) supplemented with 10% fetal bovine serum (FBS), Anti-Anti (Gibco,15240062) and gentamicin (50 µg.mL^-1^). Cal-1 cells were maintained at low passage, and only cells with less than 40 passages were used in these studies. The bovine-derived B3.13 (TX2/24) H5N1 isolate (A/Cattle/Texas/063224-24-1/2024,) was obtained from pooled milk from infected dairy cattle and used for the pathogenesis and transmission studies, infection, and virus neutralization (VN) assays in this study. The avian-derived D1.1 (NY3149/25) H5N1 isolate (A/Canada goose/New York/3149/2025), isolated from a Canada goose, was used for the comparative transmission study and for the VN assay. The working virus stocks were propagated in 10-day-old embryonated chicken eggs (ECEs) and titrated in MDCK cells using end point dilution assay and expressed as log_10_ TCID_50_.mL^-1^. Low passage working virus stocks (p. 3) were sequenced by next-generation sequencing and no mutation was found when compared with sequences of original clinical samples (GISAID accession numbers; EPI_ISL_19155861, EPI_ISL_20276734, respectively).

### Growth curves

Feline lung cells, A549, DF-1 and Cal-1 cells (2.5x10^5^ cells per well) were seeded in 12-well plates and incubated for 24 h. Cells were infected with either B31.13 (TX2/24) or D1.1 (NY3149/25) viruses at a multiplicity of infection (MOI) of 0.1 and incubated at 4°C for 1 h to allow viral adsorption. After removing the inoculum, cells were washed once with plain DMEM, and 1 mL of complete growth medium 10% FBS was added. Cells were incubated at 37°C with 5% CO, and both cells and culture supernatant were collected separately at 4, 8, 12, 24, 48, and 72 h post-infection and stored at -80°C. Time point 0 was an aliquot of virus inoculum stored at -80°C as soon as inoculation was completed. Viral titers in supernatants and cell lysates were determined in MDCK cells using endpoint dilution assays, with virus infectivity being determined by immunofluorescence assay^37,38^. Titers were calculated using the Spearman and Karber’s method and expressed as log_10_ TCID_50_.mL^-1^.

### Neuraminidase activity assay

To standardize the neuraminidase activity (NA) assay (NA-Fluor™, Applied Biosystems), we titrated the H5N1 B3.13 (TX2/24) and D1.1 (NY3149/25) virus stocks to determine the optimal virus dilution for the assay. Virus stocks were diluted in 1X assay buffer to a baseline concentration of 5 x 10^7^ PFU.mL^-1^ to enable two-fold serial dilutions. The assay was performed in triplicate using black, non-binding 96-well plates, with no virus control wells included in each plate. Serial two-fold dilutions of the viruses were prepared in 1X NA-Fluor assay buffer. The NA-Fluor substrate was reconstituted to a 2.5 mM stock solution and stored at -20°C until further use. A 200 µM working solution of the NA-Fluor substrate was prepared in 1X assay buffer prior to use and added to the virus wells and the plate was incubated at 37 °C for 60 min. The reaction was terminated by adding NA-Fluor stop solution.

Fluorescence was measured using excitation at 350–365 nm and emission at 440–460 nm and the relative fluorescence unit (RFU) values from negative control wells were averaged and subtracted from virus-containing wells to obtain background-normalized RFU values. The NA activity was plotted as RFU against each dilution of H5N1 B3.13 (TX2/24) and D1.1 (NY3149/25) viruses.

### Replicon assays

We performed replicon assays to evaluate the impact of amino acid variations in the influenza viral polymerase proteins on viral genome replication and gene transcription. We cloned viral polymerase proteins (PB2, PB1, PA) and NP of the bovine-derived B3.13 (TX2/24) and avian-derived D1.1 (NY3149/25) viruses into pCAGGS vector under the control of the chicken β-actin promoter. We chemically synthesized (GenScript, USA) a minigenome pUC57 plasmid encoding an influenza viral RNA-like segment expressing nanoluciferase (Nluc) flanked by the B3.13 TX2/24 NP segment non-coding region (NCR). Of note, the NP segment of B3.13 (TX2/24), and D1.1 (NY3149/25) are identical. The transcription of the vRNA minigenome encoding the NLuc is driven by either a human RNA polymerase I (hPolI) promoter, a cat RNA polymerase I (catPolI) promoter or a chicken RNA polymerase I (ckPolI) promoter placed upstream of the NP(NCR)-NLuc cassettes. To this end, HEK293T cells, DF-1 and feline lung cells were seeded in 24-well plate (1.25x10^5^/well) and co-transfected using PEI Prime (Sigma-Aldrich, USA) with 100 ng of each pCAGGS plasmids encoding the viral polymerase proteins (PB2, PB1, PA) and NP, together with 100 ng of the pUC57-NP(NCR)-NLuc minigenome plasmid. Cells transfected with all influenza RNP proteins except the PB1-encoding plasmid served as a negative control. For luciferase assays, cell culture supernatant was removed after 24 and 48 hours and cells were lysed with 50 µL of 1X Passive Lysis Buffer (Promega), and Nluc expression was quantified using a Nano-Glo luciferase assay system (Promega, USA).

### Pathogenesis of HPAI H5N1 in cats

To investigate the pathogenesis, infection dynamics and tissue tropism of H5N1B3.13 virus in cats, twelve (n = 12) H5N1-seronegative research cats of 15-18 months age obtained from Marshall Bioresources and housed individually in Horsfall HEPA-filtered cages in the animal biosafety level 3 (ABSL-3) facility at the East Campus Research Facility (ECRF) at College of Veterinary Medicine, Cornell University. Food and water were provided *ad libitum*. After acclimation, nasal swabs and blood samples were collected from all animals and tested by rRT-PCR for influenza A virus and neutralizing antibodies against H5N1 virus. On day 0, nine (n = 9) cats were anesthetized and inoculated oronasally with 2 mL virus suspension (1 mL orally, 1 mL intranasally) of a H5N1 clade 2.3.4.4b genotype B3.13 virus A/Cattle/Texas/06322424-1/2024, (EPI_ISL_19155861)^10^ containing 1x10^4.5^ PFU.mL^-1^. Three (n = 3) control cats were mock inoculated with 2mL minimal essential media (MEM) through the same routes. Following inoculation, animals were monitored daily for clinical signs consistent with influenza A H5N1 virus infection. Clinical disease was assessed using a clinical score index (CSI) from days 1 to 7 post-inoculation (pi) as previously described.^39^ Briefly, the CSI was calculated based on daily individual observations and included five predefined clinical parameters; hunched posture (3 points), ruffled fur (3 points), reduced food or water intake (defined as consumption of <25% of provided food and water; 2 points), body weight loss >15% (10 points), and neurological signs (e.g., hind-limb paralysis, tremors; 10 points). The presence of the above clinical signs led to assignment of the full score for that parameter, whereas the absence of clinical signs was scored as 0, ensuring objectivity and stringency of the clinical assessment. Cats with CSI >15 were subjected to increased monitoring frequency (2X per day). Rectal body temperature and body weight were measured and recorded daily throughout the challenge period. A body weight loss ≥20% was defined as the humane endpoint for euthanasia and necropsy. Nasal (NS), oropharyngeal (OPS), and rectal (RS) swabs were collected on days 0, 1, 3, 5, and 7 pi in 1 mL of virus transport medium (VTM) and stored at −80 °C until further analysis. Whole blood samples were collected on days 3, 5, 6, and 7 pi in EDTA-coated tubes and stored at −80 °C and at 4 °C. Serum samples were collected in serum separation tubes on days 3, 5, 6, and 7 pi. On scheduled necropsy days (3, 5, 7 pi), all cats were euthanized in accordance with the American Veterinary Medical Association (AVMA) Guidelines for the Euthanasia of Animals. Briefly, cats were sedated via intramuscular administration of dexmedetomidine (Dexdomitor®, 0.01–0.04 mg kg ¹), followed by induction of deep anesthesia using inhaled isoflurane (3–5%). The absence of the pedal withdrawal reflex confirmed adequate depth of anesthesia. Once deep anesthesia and unconsciousness were verified, blood was collected via the intracardiac route until respiratory arrest occurred. Cats were monitored for a minimum of 3 minutes to confirm death before necropsy. For histopathological evaluations, we collected multiple tissues, including nasal turbinates, trachea, lung, heart, liver, spleen, kidney, stomach, pancreas, intestine, testes/ovary, uterus, mammary gland, brain, tonsils, retropharyngeal lymph nodes (RPLN), mediastinal LN, and mesenteric LN. Tissues were fixed in 10% neutral buffered formaldehyde for 48 hours and then replaced with 80% ethanol to be stored at room temperature until processing.

### Transmission efficiency of H5N1 B3.13 and D1.1 genotypes in cats

To assess the transmission efficiency of H5N1 B3.13 and D1.1 genotypes, a total of sixteen (n = 16) research cats (kindly donated by Dr. Jorge Osorio at the University of Wisconsin-Maddison) were allocated into two groups: H5N1 B3.13 (TX2/24) (n = 8), and H5N1 D1.1 (NY3149/25) (n = 8). Each group was further divided into infected (I, n = 4) and contact (C, n = 4) subgroups. Each animal in the infected subgroup was inoculated oronasally (1 mL orally and 1 mL intranasally) with 1 X 10^4.5^ PFU.mL^-1^ of B3.13 (TX2/24)^10^ or D1.1 (NY3149/25) viruses. At 24 hours post-inoculation (pi), the infected cats were transferred to the enclosure of the contact animals, paired and cohoused (1:1) with the contact animals for up to 14 days. Nasal (NS), oropharyngeal (OPS), and rectal (RS) swabs and serum as well as environmental enclosure swabs (floor, feeder, and water), were collected daily to quantify viral RNA and infectious virus by rRT-PCR and virus titrations, respectively. Throughout the study period (14 days), cats reaching the humane endpoint criteria were euthanized. All remaining cats were euthanized on day 14 pi, and blood and tissue samples were collected as above.

### RNA extraction and rRT-PCR

Viral RNA was extracted from swab samples and tissue homogenates (10%, w/v) using the IndiMag Pathogen kit (Indical Bioscience SP947855P196, Germany) in an automated KingFisher flex (ThermoFisher) extractor. Real-time reverse transcriptase PCR (rRT-PCR) was performed on purified RNA using the Path-ID Multiplex One-Step RT-PCR kit (ThermoFisher, USA) and influenza A matrix gene-specific primers and probe. Both negative and positive controls were run along with the test samples. A standard curve using serial 10-fold dilution of H5N1 B3.13 (TX2/24) viral RNA was performed to estimate viral RNA copy number (log_10_ genome copies.mL^-1^) using the relative quantification method, and values were plotted in GraphPad Prism Prime Software version 10.1.2 (GraphPad, La Jolla, CA, USA).

### Virus titrations

The rRT-PCR positive samples collected during pathogenesis studies in cats were titrated in embryonated chicken eggs (ECE). To determine the embryo infectious dose 50 (EID_50_.mL^-1^), 10-day-old ECEs were inoculated with 100 μL of 10-fold serial dilutions of swab sample in triplicate via the allantoic cavity route and incubated at 37.6 °C and 50% relative humidity. The eggs were candled daily, and the embryo mortality was recorded. Embryos that survived after 96 h were chilled. Allantoic fluid was collected and tested for hemagglutination activity using 0.5% turkey red blood cells. Virus titers were calculated using Reed and Munch method and expressed as log_10_ EID_50_.mL^-1^.

The rRT-PCR positive samples collected from the transmission study in cats were titrated in MDCK cells using the endpoint dilution method. For that MDCK cells were seeded in 96-well plates at 2.5 x 10^4^ per well, and 24 h post-seeding, the plates were moved to BSL3. Serial 10-fold dilutions of samples were prepared in DMEM and inoculated in quadruplicate (50 μL/well) into MDCK cells. After 1 h of incubation at 37 °C, 50 μL of complete growth media, DMEM supplemented with 5% FBS, was added to each well of the 96-well plate. After 48 h of incubation at 37 °C, cells were fixed with 3.7% formaldehyde for 30 min at room temperature. Infected cells were detected by immunofluorescence assay (IFA) using the mouse anti-NP mAb (HB65) mouse monoclonal antibody followed by goat anti-mouse IgG (H&L) conjugated with Alexa Fluor 594 (ImmunoReagents Laboratories) secondary antibody incubation. Virus titers were calculated using Spearman and Karber’s method and expressed as log_10_ TCID_50_.mL^-1^.

### Virus neutralization assay

The H5N1 B3.13 (TX2/24) and D1.1 (NY3149/25) viruses were used to assess seroconversion of the cats during pathogenesis and transmission studies as described previously^40^. Briefly, 75 μL MEM was added to the first column (A1 to H1) of a 96-well plate and 50 µL MEM was added to the remaining wells. Heat-inactivated serum (25 µL) was added to the first well, mixed 5 times with a pipette, and 50 μL was transferred to the second well. A serial two-fold dilution was prepared (1:8 to 1:16384). A virus suspension containing 200 TCID_50_.mL^-1^ in 50 µL MEM was added to all wells and the virus-serum mixture was incubated at 37 °C for 1 h. Known positive and negative control serum samples were used in each run. After that 100 μL of Cal-1 cells (approximately 2.5 x10^4^ per well) were added to each well. After 48 h of incubation at 37 °C, cells were fixed with 3.7% formaldehyde, and IFA using an anti-influenza A NP antibody was performed as described above. Neutralizing antibody titers were determined as the reciprocal of the highest serum dilution to completely neutralize H5N1 virus replication, as evidenced by the absence of fluorescence.

### Histopathology and *in situ* hybridization

Formalin-fixed tissues were processed, paraffin-embedded and sectioned at 5 µm thickness and stained with hematoxylin and eosin (H&E). Histological changes were examined by two board-certified anatomic pathologists blinded to the experimental details. For all tissue, the degree of inflammation, necrosis, and hemorrhage was scored on the entirety of the section area as follows 0: none, 1: mild (affecting <25% of the tissue), 2: moderate (affecting >25% and less than 50% of the tissue), 3: severe (affecting >50% of the tissue). To determine virus tropism and tissue distribution, we performed RNA-Scope *in situ* hybridization (ISH) on tissue sections as previously described^41^. Briefly, tissue sections were deparaffinized with xylene, dehydrated with absolute ethanol and blocked with peroxidase. After antigen retrieval for 1 hour, V-InfluenzaA-H5N8-M2N1 probe (Advanced Cell Diagnostics) and the RNA-Scope HD2.5 assay were used as per the manufacturer’s instructions.^40^ All the slides were scanned at 40X resolution, and digital slides were examined for virus tropism and tissue distribution.

### Fluorescent in situ hybridization (FISH)

Brain tissues were examined for glial cell tropism of H5N1 B3.13 virus using RNAscope™ Multiplex Fluorescent V2 kit (UM 323100, ACDBio). To this end, we custom-designed *Felis catus* specific probes targeting vascular endothelial cells (CLDN5; Claudin-5), neurons (MAP2; microtubule-associated protein-2), and astrocytes (GFAP; Glial Fibrillary Acidic Protein). A HPAI influenza virus marker (V-Influenza A-H5N8-M2M1-C1) was used to colocalize viral RNA in tissues. Briefly, 5µm thick tissue sections were deparaffinized with xylene and washed with ethanol and distilled water. After hydrogen peroxide treatment for 10 minutes at room temperature, antigen unmasking was performed by boiling the slide in RNAscope 1X Target Retrieval Reagent for 20 minutes. Following the protease treatment at 40 °C for 30 minutes, tissues were sequentially probed. First, tissues were hybridized with viral V-Influenza A-H5N8-M2M1-C1 probe for 2 hours at 40 °C, followed by three rounds of amplification with amplifying probes for 30 minutes each at 40 °C. To label the probe 1(V-Influenza A-H5N8-M2M1-C1) with horse reddish peroxidase (HRP), tissues were incubated with horse (HRP-C1) at 40 °C for 15 minutes. To develop signals from probe 1 (HPAI H5N1 viral probe), 200 µL of compatible Opal dye 520 fluorophore (Opal 520 Reagent Pack (FP1487001KT) was added to the tissues with probe-HRP complex and incubated for 30 minutes at 40 °C. Finally, HRP blocker was added to the slides to prepare for subsequent probes (MAP2, GFAP, and CLDN5). We used Opal 570 (FP1488001KT) for MAP2, Opal 650 (FP1496001KT) for GFAP, and Opal 780 (FP1501001KT) for CLDN5 (Supplementary table 1).

### Statistical analysis

Statistical analysis was performed by two-way analysis of variance (ANOVA) followed by multiple comparisons and mean, and standard error of mean values were presented. GraphPad Prism Prime (version 15.1) was used to plot the data.

## Data availability

All data supporting the findings of this study are available within the manuscript, supplemental figures, and supplementary table files.

## Acknowledgement

We would like to thank Cornell EH&S, Biosafety, and CARE teams for their help in setting up the protocols and procedures to conduct animal work with HPAI H5N1 virus at the Cornell ABSL-3 facilities. The research cats used in the transmission studies were kindly donated by Dr. Jorge Osorio from the University of Wisconsin-Madison. Figures 1A, 3C, 4A, 4C, 6, and Figure S2A, were prepared with Biorender.

## Author contributions

D.G.D. designed and supervised the study. S.L.B., R.R., M.N., and P.B.S.O. conducted the animal studies. S.L.B., M.N., and D.G.D. analyzed the experimental data. E.A.D. and G.R.H. reviewed and scored the in situ hybridization and histological lesions. S.L.B., M.N. and D.G.D. drafted the manuscript with input from all authors.

## Declaration of generative AI and AI-assisted technologies in the manuscript preparation process

During the preparation of this work the author(s) used Claude (Anthropic) to improve grammar and readability of this manuscript. Figure S2A was created by inputting the original brain tissue scan from day 3 into BioRender AI tool to depict different anatomic regions of the brain. After using this tool/service, the author(s) reviewed and edited the content as needed and take(s) full responsibility for the content of the published article.

---

**Figure S1:**
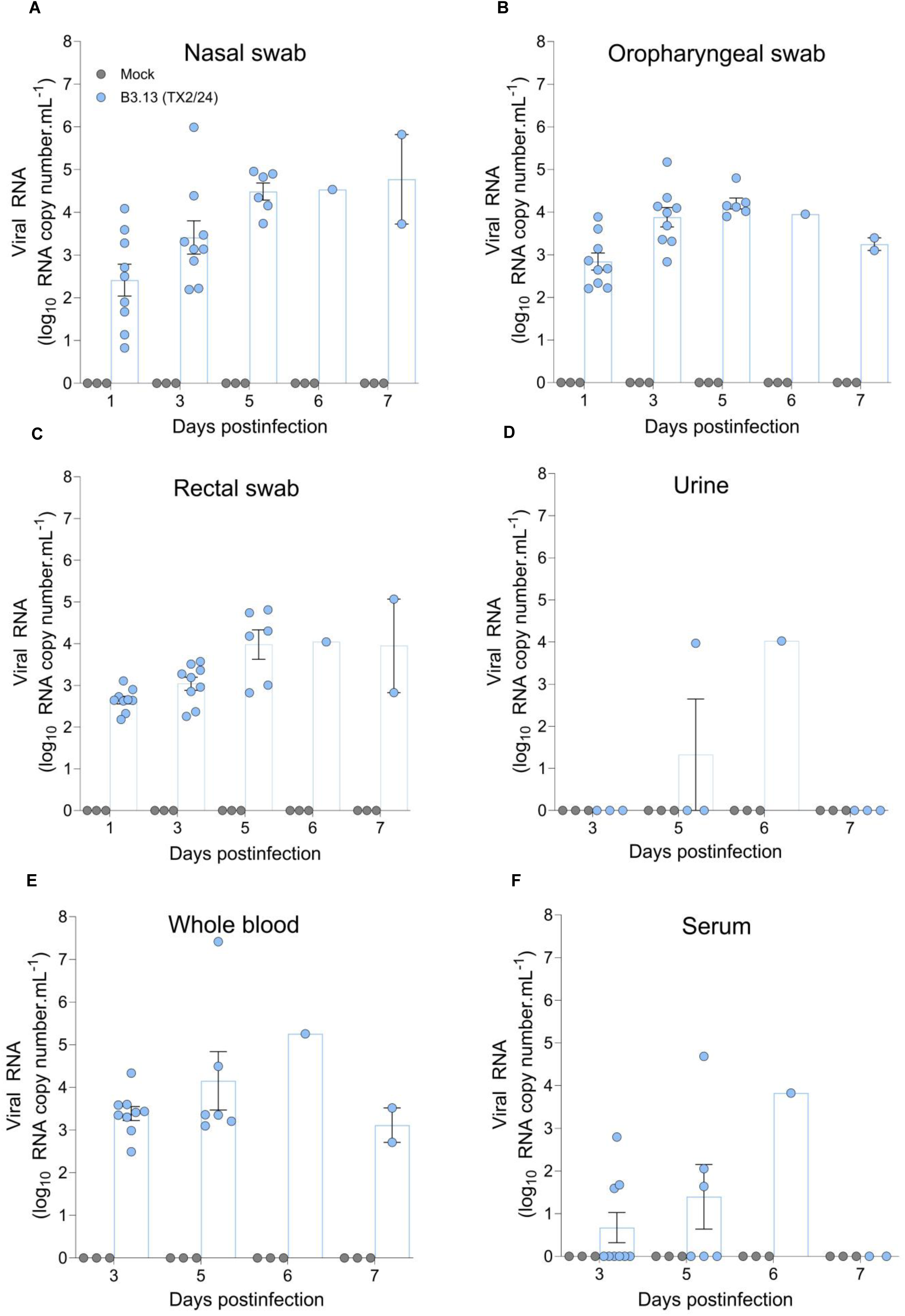
H5N1 viral RNA shedding and viremia detected in natural secretions . Viral RNA from mock and infected cat quantified by qPCR in nasal **(A)**, oropharyngeal **(B)**, and rectal swabs **(C)**, urine **(D)**, whole blood **(E)**, and serum **(F)**. Each dot in the bar graphs indicates an individual animal from each group (mean with SEM, n = 3 cats/mock group, 9 cats/ H5N1 (TX2/24) infected group). Related to Figure 1.

**Figure S2:**
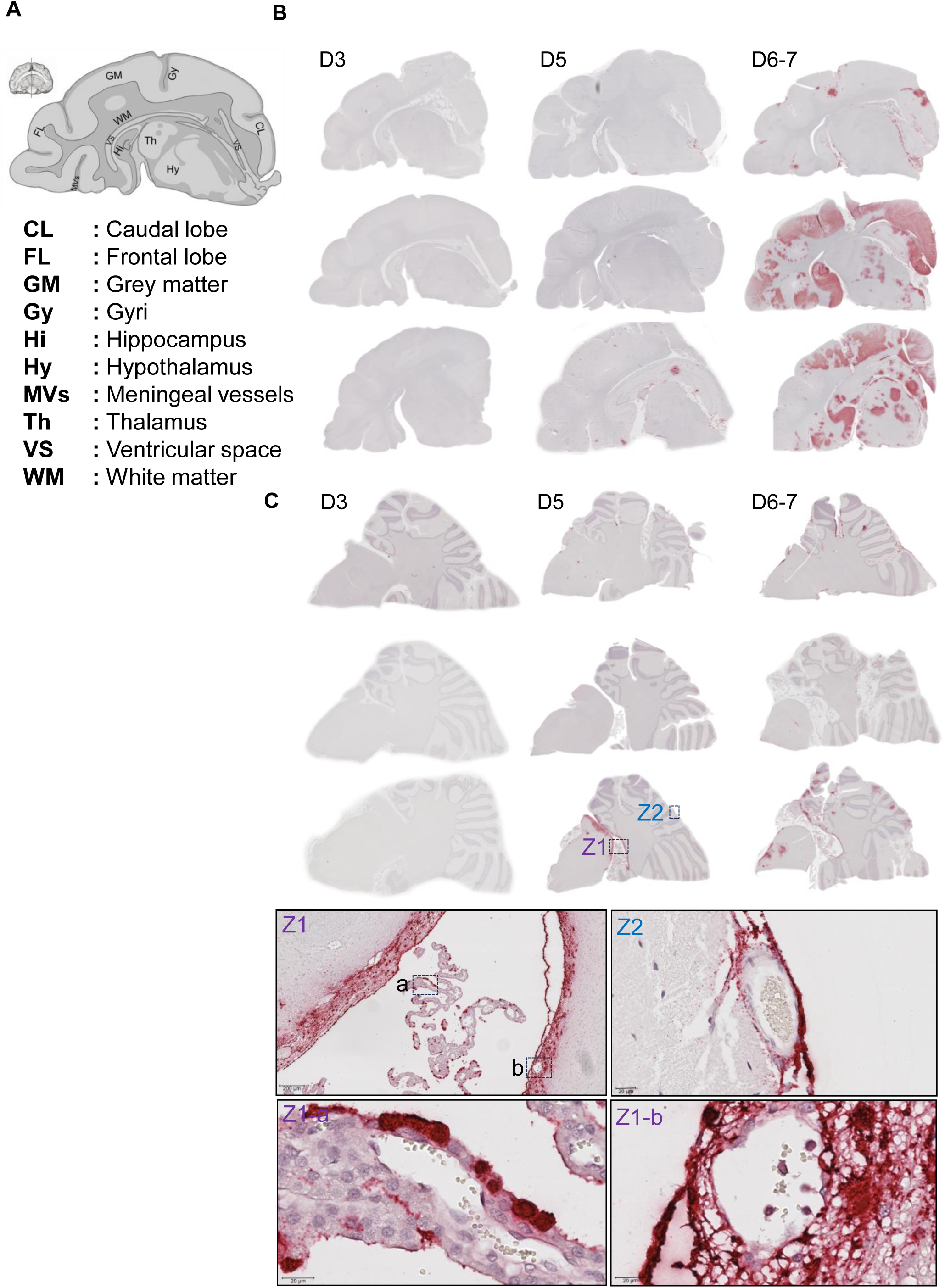
Neuroinvasion dynamics of H5N1 B3.13 virus in cat brain tissue. **A)** A schematic diagram of the cat brain depicting different anatomical regions of the brain. Temporal spread of H5N1 viral RNA demonstrated in cerebrum **(B)** and cerebellar **(C)** tissues. The higher magnification (Z labelled areas) shows viral RNA labeling in ependymal cells (Z1a) and meningeal blood vessels (Z1b). Related to Figure 2.

**Figure S3:**
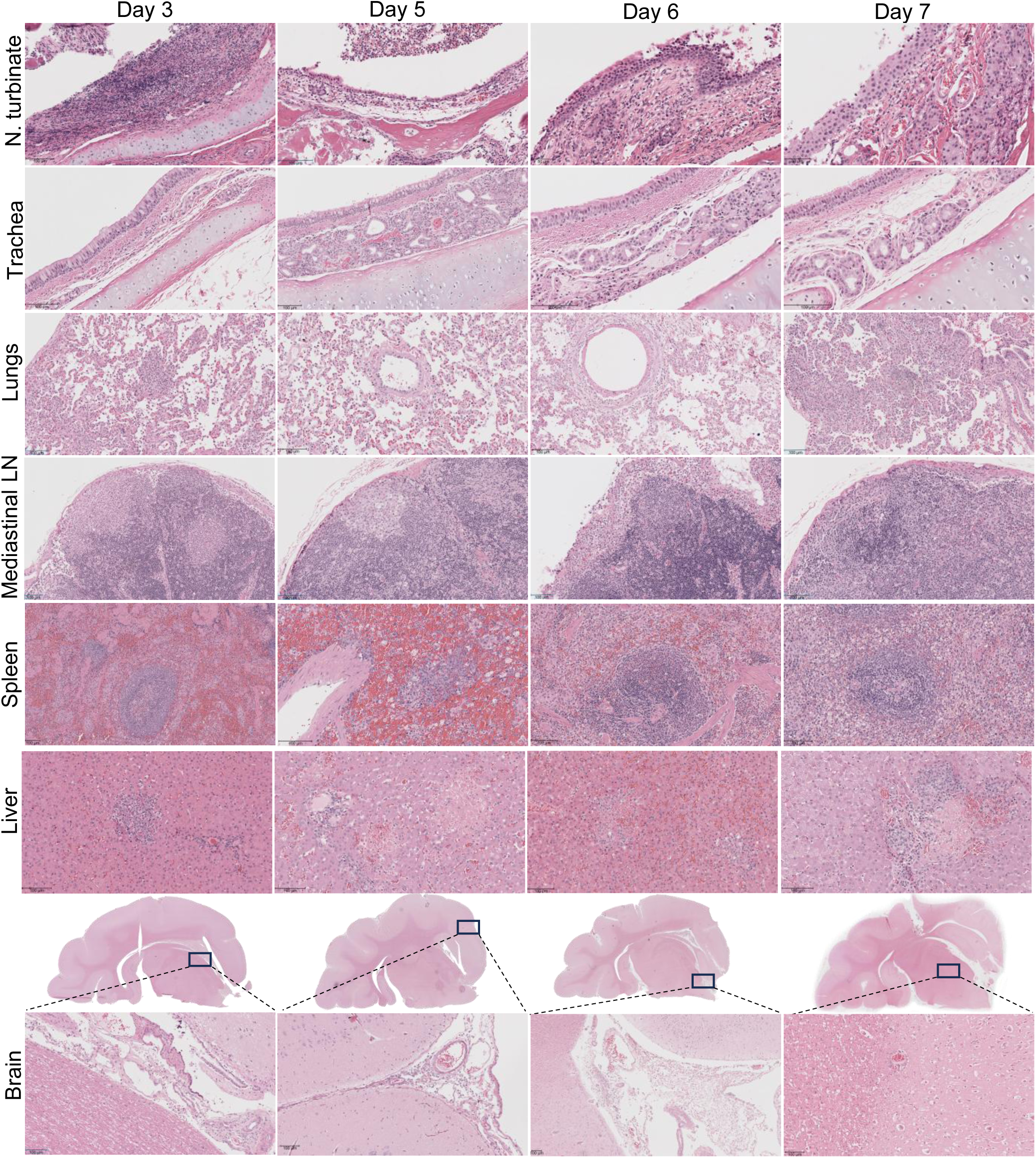
Histopathological lesions of HPAI H5N1 in different tissues. Hematoxylin and eosin staining show histological changes in nasal turbinates, trachea, lung, mediastinal lymph node, spleen, liver and brain on 3-, 5-, 6- and 7-day post-infections. Related to Figure 2.

**Figure S4:**
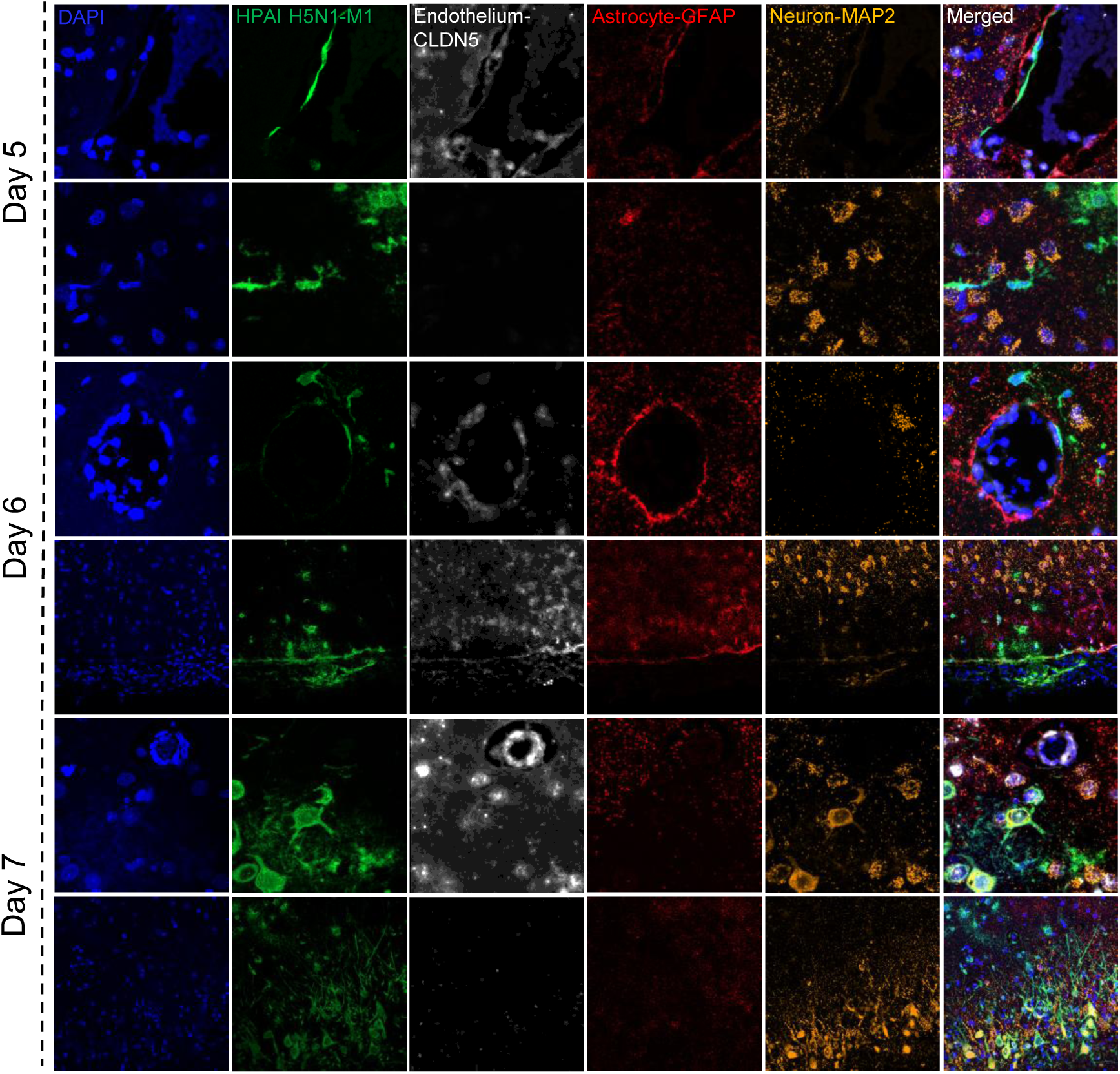
Neurotropism of HPAI H5N1 B3.13 in cat brain on 5-, 6-, 7-days post infection. Fluorescent in situ hybridization (FISH) showing HPAI H5N1 virus (green) infection in vascular endothelial cells (grey), astrocytes (red), and neurons (orange). Blue is DAPI-stained cellular nuclei. Related to Figure 3.

**Figure S5:**
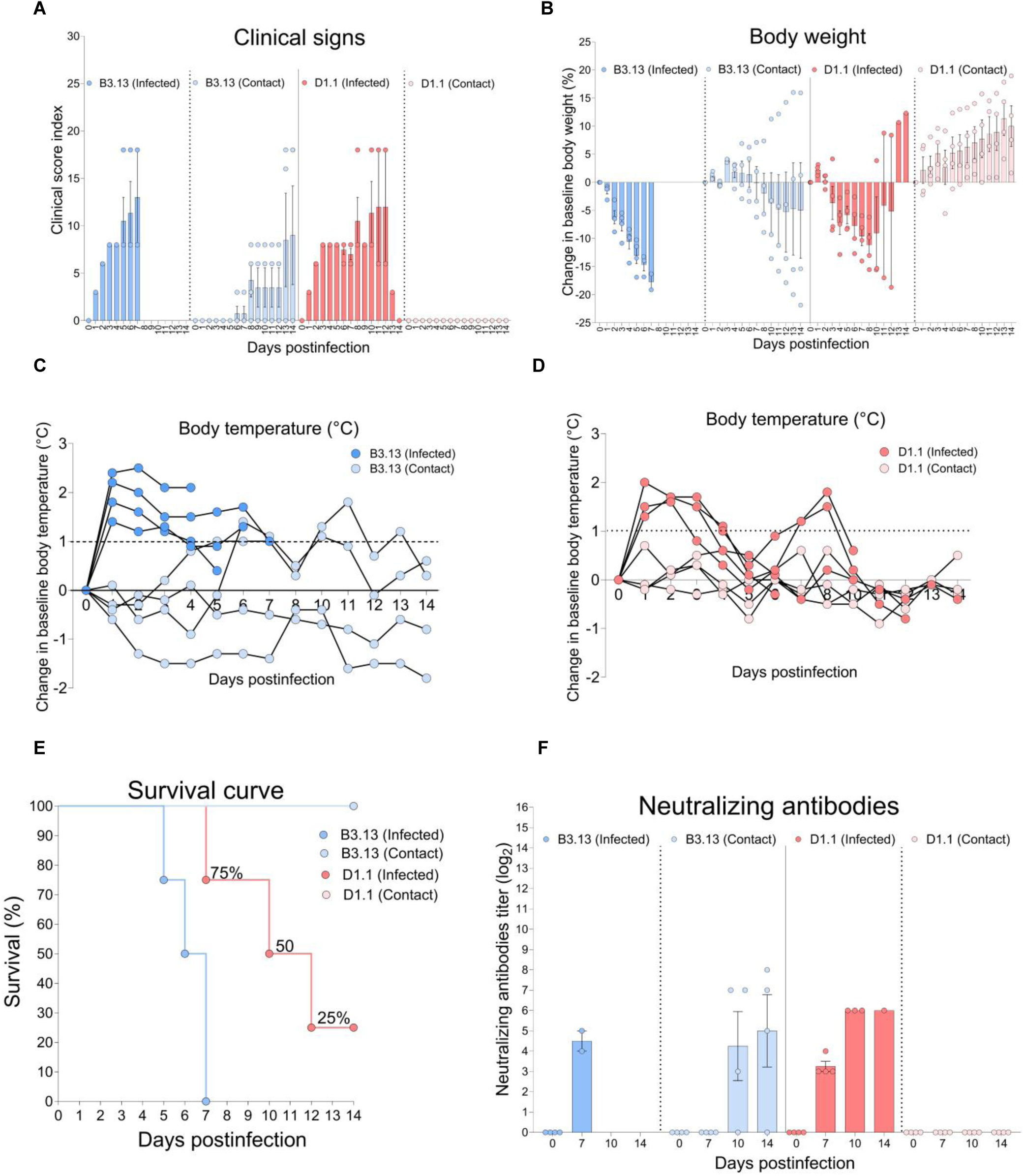
Clinical outcome of HPAI H5N1 B3.13 and D1.1 genotype infection in cats. **A)** Clinical scores of indexed (infected) and direct contact cats **B)** Percent change in baseline body weight of indexed and direct contact cats **C-D)** change in baseline body temperature of indexed and direct contact cats. **E)** Percent survival of indexed and direct contact cats. **F)** Seroconversion of indexed and direct contact cats. Related to Figure 4.

**Figure S6:**
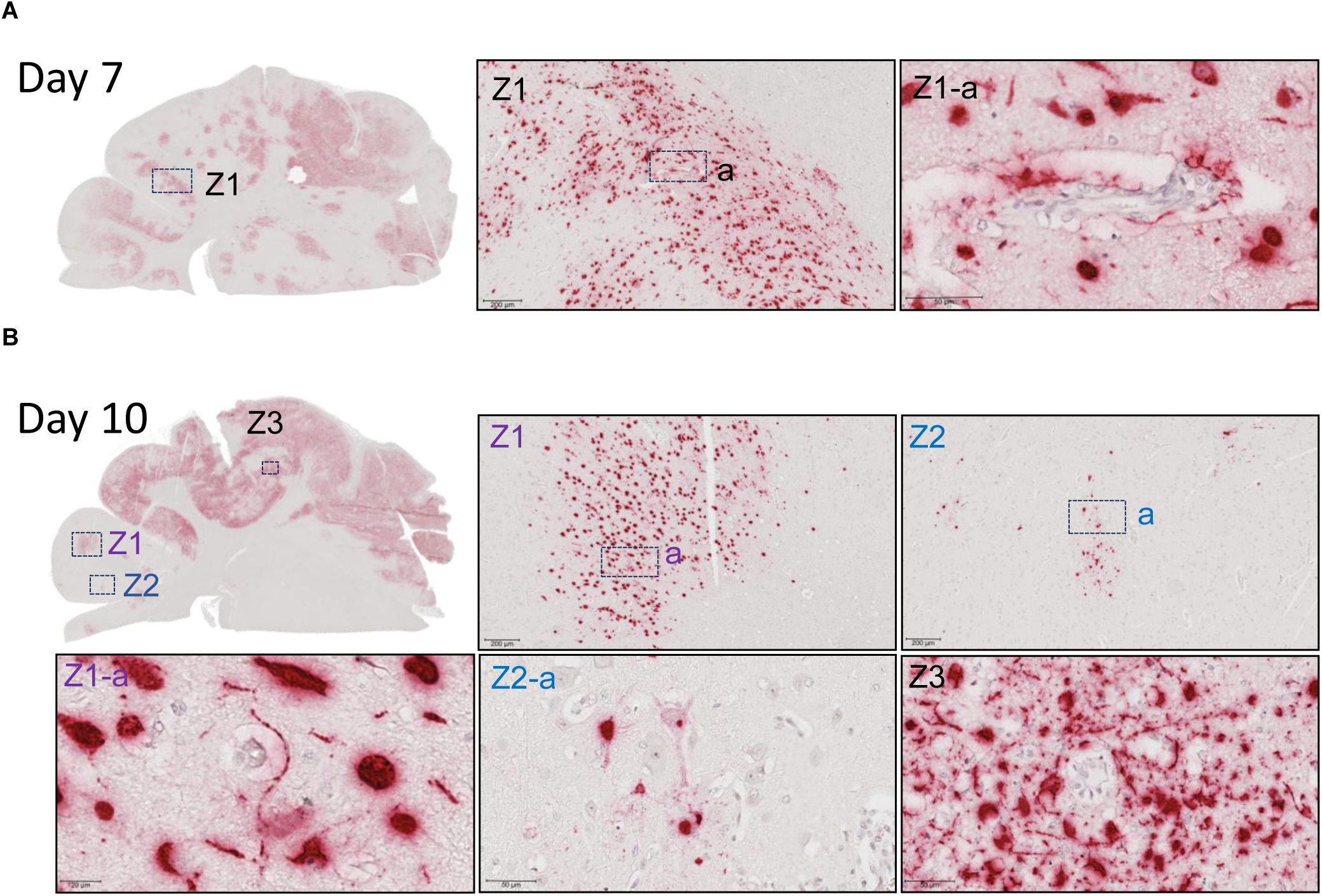
Temporal spread of HPAI H5N1 D1.1 genotype in brain tissue. In situ hybridization showing HPAI H5N1 D1.1 genotype viral RNA in infected cats at days 7 and 10 post-inoculation. **A)** The higher magnification of Z-labelled areas (Z1a) shows viral RNA labeling of blood vessels and cells in perivascular areas at day 7 post-inoculation. **B)** The high magnification of Z-labelled areas (Z2a) shows viral RNA labeling in cells of the perivascular area and likely transport of virus to the connected neurons. Related to Figure 3.

**Figure S7:**
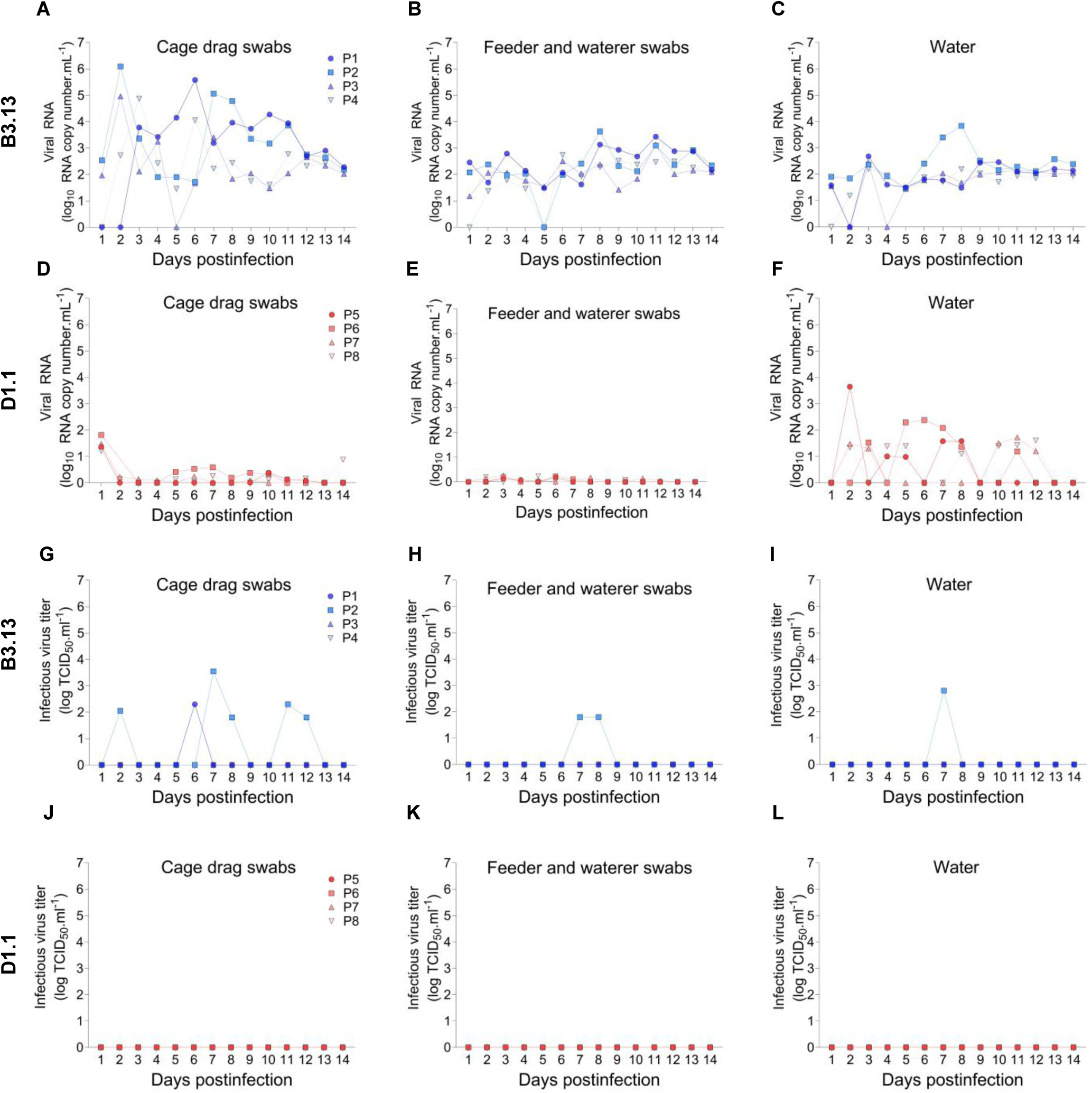
HPAI H5N1 B3.13 and D1.1 genotypes in environmental samples. Viral RNA loads (log₁₀ copies mL⁻¹) were quantified in **(A, D)** cage drag swabs, **(B, E)** feeder and waterer swabs, and **(C, F)** water samples collected from cat pairs exposed to the B3.13 or D1.1 genotypes, respectively. Infectious virus titers (TCID₅₀ mL⁻¹) were determined in corresponding environmental samples, including **(G, J)** cage drag swabs, **(H, K)** feeder and waterer swabs, and **(I, L)** water samples from B3.13- and D1.1-exposed cat pairs. Related to Figure 4.

