## Supplementary Table 1 for "Hematogenous neuroinvasion and genotype-dependent transmission of influenza A H5N1 viruses in the cat host"

**Table S1.** Details of probes and Opal dyes used in 4-plex fluorescent in situ hybridization (FISH) assay

| **Target** | **Probes** | **Opal dye** | **Probe mix ratio** | **Opal dye dilution** |
|---|---|---|---|---|
| HPAI H5N1 | V-Influenza A-H5N8-M2M1-C1 | Opal 520 | 1:50 | 1:300 in TSA buffer |
| Neuron | MAP2 | Opal 570 | 1:50 | 1:300 in TSA buffer |
| Astrocyte | GFAP | Opal 650 | 1:50 | 1:300 in TSA buffer |
| Vascular endothelial cells | CLDN5 | Opal 780 | 1:50 | 1:300 in TSA buffer |

- The 4-plex positive control is a RTU mixture of four probes targeting common housekeeping genes POLR2A in channel C1, PPIB in channel C2, UBC in channel C3, and HPRT in channel 4.

- The 3-plex/4-plex negative control probes are RTU mixture of 3/4 probes targeting dapB which is a soil bacterium gene. Each detection channel has its own negative control probe: dapB-C1, dapB-C2, dapB-C3 and/or dapB-C4
