## Supplementary Table 2 for "Hematogenous neuroinvasion and genotype-dependent transmission of influenza A H5N1 viruses in the cat host"

**Table S2. S**ubstitution mutations across genome fragments of HPAI H5N1 genotypes B3.13 (TX2/24) and D1.1 (NY3149/25)

## PB2

| GISAID Accession # | Isolate | T58A | G79S | V109I | V139I | N195D | E362G | D441N | I444V | V478I | V495I | M631L | V649I | T676A | S715N |
|---|---|---|---|---|---|---|---|---|---|---|---|---|---|---|---|
| EPI_ISL_19155861 | B3.13 (TX2/24) | A | S | I | I | D | G | N | V | I | I | L | I | A | N |
| EPI_ISL_20276734 | D1.1 (NY3149/25) | | S | | | D | | | V | I | | | | | N |
| EPI_ISL_19634827 | D1.1 (LA/12/24) | | S | | | D | | | V | I | | | | | N |

## PB1

| GISAID Accession # | Isolate | N16D | T59S | E75D | G104E | R135K | G154S | M171V | D172E | T176K | M179I | I191V | K207R | I372M | N375S | R430K | K571R | A587P | E614D | E618D | D619E | I660V | S694N | G738E |
|---|---|---|---|---|---|---|---|---|---|---|---|---|---|---|---|---|---|---|---|---|---|---|---|---|
| EPI_ISL_19155861 | B3.13 (TX2/24) | | S | D | E | | | V | E | K | I | V | | M | S | K | R | P | | | | V | N | E |
| EPI_ISL_20276734 | D1.1 (NY3149/25) | D | | | E | K | S | V | | K | | V | R | M | | K | R | | D | D | | V | | E |
| EPI_ISL_19634827 | D1.1 (LA/12/24) | D | | | E | | S | | | K | | V | R | M | | K | R | | D | | E | V | | E |

## PA

| GISAID Accession # | Isolate | M61I | A85T | K113R | M211I | L219I | R269K | S277P | I322V | V323I | I348L | S388G | R391K | E399V | S400P | V441M | C489S | K497R | E538G | I545V | S558L | S608T | K626R |
|---|---|---|---|---|---|---|---|---|---|---|---|---|---|---|---|---|---|---|---|---|---|---|---|
| EPI_ISL_19155861 | B3.13 (TX2/24) | | | R | | I | | P | | | | | | | | | | R | | | L | | |
| EPI_ISL_20276734 | D1.1 (NY3149/25) | I | T | | | | K | | V | I | L | G | K | | P | M | | | | V | | T | R |
| EPI_ISL_19634827 | D1.1 (LA/12/24) | I | T | | I | | K | | V | I | L | G | K | V | P | M | S | | G | V | | T | R |

## HA

| GISAID Accession # | Isolate | V11I | T52A | M120L | L131Q | T211I | A226V | K341R | N491D | V526I |
|---|---|---|---|---|---|---|---|---|---|---|
| EPI_ISL_19155861 | B3.13 (TX2/24) | | | | Q | I | | | | |
| EPI_ISL_20276734 | D1.1 (NY3149/25) | I | A | L | | | V | R | D | I |
| EPI_ISL_19634827 | D1.1 (LA/12/24) | I | A | L | | | V | R | D | I |

## NA

| GISAID Accession # | Isolate | T8I | V20I | M23V | Y44N | Q45H | P48T | I53V | V67I | F74L | L75I | T81D | S82P | S82T | T84A | V149I | N221S | V234I | V241I | K257R | L269M | G286S | D287E | I288V | V321I |
|---|---|---|---|---|---|---|---|---|---|---|---|---|---|---|---|---|---|---|---|---|---|---|---|---|---|
| EPI_ISL_19155861 | B3.13 (TX2/24) | | | | | | | | I | | | | | | | | | | | | M | | | | I |
| EPI_ISL_20276734 | D1.1 (NY3149/25) | I | I | V | N | H | T | V | I | L | I | D | | T | A | I | S | I | I | R | | S | E | V | |
| EPI_ISL_19634827 | D1.1 (LA/12/24) | I | I | V | N | H | T | V | I | L | I | D | P | | A | | S | I | I | R | | S | E | V | |

## NA

| GISAID Accession # | Isolate | N329S | S336G | M338V | S339P | E395A | T397M | M418I |
|---|---|---|---|---|---|---|---|---|
| EPI_ISL_19155861 | B3.13 (TX2/24) | | | | P | | | |
| EPI_ISL_20276734 | D1.1 (NY3149/25) | S | G | V | | A | M | I |
| EPI_ISL_19634827 | D1.1 (LA/12/24) | S | G | V | | A | | |

## NS

| GISAID Accession # | Isolate | S7L | G53D | E75G | T76A | P83S | S87P | V111A | C116S | D139N | L147I | D171N | R193Q | A223E |
|---|---|---|---|---|---|---|---|---|---|---|---|---|---|---|
| EPI_ISL_19155861 | B3.13 (TX2/24) | L | D | | A | S | P | | S | N | | | | E |
| EPI_ISL_20276734 | D1.1 (NY3149/25) | | D | G | A | | | | | N | I | N | Q | |
| EPI_ISL_19634827 | D1.1 (LA/12/24) | | D | G | A | | | A | | | I | N | Q | |

## NP

| GISAID Accession # | Isolate | R50S | H52Y | V105M | P318L | I353V | S450N | S482N |
|---|---|---|---|---|---|---|---|---|
| EPI_ISL_19155861 | B3.13 (TX2/24) | S | | M | | V | N | N |
| EPI_ISL_20276734 | D1.1 (NY3149/25) | S | Y | M | | V | N | |
| EPI_ISL_19634827 | D1.1 (LA/12/24) | S | Y | M | L | V | N | |

## MP

| GISAID Accession # | Isolate | N82S | N85S | N87T | V200A | A227T |
|---|---|---|---|---|---|---|
| EPI_ISL_19155861 | B3.13 (TX2/24) | S | S | T | | T |
| EPI_ISL_20276734 | D1.1 (NY3149/25) | | | | A | |
| EPI_ISL_19634827 | D1.1 (LA/12/24) | | | | A | |
